# RUV-III-NB: Normalization of single cell RNA-seq Data

**DOI:** 10.1101/2021.11.06.467575

**Authors:** Agus Salim, Ramyar Molania, Jianan Wang, Alysha De Livera, Rachel Thijssen, Terence P. Speed

**Affiliations:** Melbourne School of Population and Global Health, University of Melbourne VIC 3053; Bioinformatics Division, Walter and Eliza Hall Institute of Medical Research, Parkville VIC 3052; School of Mathematics and Statistics, University of Melbourne VIC 3010; Baker Heart and Diabetes Institute, Melbourne VIC 3004; Department of Mathematics and Statistics, La Trobe University VIC 3086; Department of Medical Biology, University of Melbourne VIC 3010; School of Science, RMIT University, Melbourne VIC 3000; Blood Cells and Blood Cancer Division, Walter and Eliza Hall Institute of Medical Research, Parkville VIC 3052, Australia

## Abstract

Despite numerous methodological advances, the normalization of single cell RNA-seq (scRNA-seq) data remains a challenging task and the performance of different methods can vary greatly across datasets. Part of the reason for this is the different kinds of unwanted variation, including library size, batch and cell cycle effects, and the association of these with the biology embodied in the cells. A normalization method that does not explicitly take into account cell biology risks removing some of the signal of interest. Furthermore, most normalization methods remove the effects of unwanted variation for the cell *embedding* used for clustering-based analysis but not from gene-level data typically used for differential expression (DE) analysis to identify marker genes. Here we propose RUV-III-NB, a statistical method that can be used to remove unwanted variation from both the cell *embedding* and gene-level counts. RUV-III-NB explicitly takes into account its potential association with biology when removing unwanted variation via the use of pseudo-replicates. The method can be used for both UMI or sequence read counts and returns adjusted counts that can be used for downstream analyses such as clustering, DE and pseudotime analyses. Using five publicly available datasets that encompass different technological platforms, kinds of biology and levels of association between biology and unwanted variation, we show that RUV-III-NB manages to remove library size and batch effects, strengthen biological signals, improve differential expression analyses, and lead to results exhibiting greater concordance with independent datasets of the same kind. The performance of RUV-III-NB is consistent across the five datasets and is not sensitive to the number of factors assumed to contribute to the unwanted variation. It also shows promise for removing other kinds of unwanted variation such as platform effects. The method is implemented as a publicly available R package available from https://github.com/limfuxing/ruvIIInb.

## Introduction

Single-cell RNA-seq (scRNA-seq) technologies have gained popularity over the last few years as more and more studies interrogate transcriptomes at the single cell level. Just as with other omics data, scRNA-seq data inevitably contains unwanted variation which can compromise downstream analyses if left unaddressed. As in the case with bulk RNA-seq data, library size is the major source of unwanted variation in scRNA-seq data and consequently, removing library size effects is the first priority in preprocessing scRNA-seq data. The successful removal of library size effects is crucial for the validity of downstream analyses such as clustering, cell-type annotation, differential expression and trajectory analyses. Several studies^1,2,3,4^ have found that the bulk RNA-seq procedures for removing library size effects do not work well for scRNA-seq data. This is because the relationship between gene expression and library size in scRNA-seq data is typically complex and gene-specific, a feature of the data that has necessitated the development of methods using gene-specific scaling factors,^3,4,5^ as opposed to methods that use global scaling factors e.g.^6,1^ In addition to library size effects, scRNA-seq data can exhibit batch effects^7^ due to variation between cell counts *within* a study (e.g. due to plate-to-plate variation) and variation between cell counts *across* studies (e.g. due to platform and sample preparation variation). In this paper, we concentrate on dealing with the first, although we show that our method has the potential to perform *data integration* by adjusting for library size and batch effects across studies.

Like Vallejos *et al*.,^2^ in this paper we will use the term ‘normalization’ to refer to a procedure that attempts to remove all kinds of unwanted variation and not only that due to library size. One of the key challenges when performing normalization is to remove the right kind and amount of variation. Removing the wrong or too much variation risks removing biology, especially if biological variation is associated with unwanted variation. Most methods that adjust scRNA-seq data for batch effects^8,9,10,11^ proceed in two steps: library size effects are removed first, and then batch effects are removed from data that has been adjusted for library size. This approach is reasonable if there is little or no association between library size, batch and biology, but when there are such associations, its effectiveness may be reduced. For example, when different cell-types have quite different library size distributions, the first step may adjust the data too aggressively and remove library size differences arising as differences between cell-types. ZINB-WaVE^12^ can be used to perform simultaneous adjustment for library size and batch effects. However, it requires that the batches are known a priori, and its adjustment is carried out without considering the possibility that library size, biology and batch may be associated. Furthermore, most normalization methods remove the effects of unwanted variation for the cell *embedding* used for clustering-based analysis but may severely distort gene-level data used for differential expression (DE) analysis used to identify marker genes.^13^

In this paper, we propose RUV-III-NB that simultaneously adjusts scRNA-seq gene counts for library size and within study batch differences. As with RUV-III^14^ which inspired this work we do not assume that batch details are known, but seek to use *replicates* and *negative control* genes to capture and adjust for the unwanted variation. Negative controls are genes whose variation is (largely) unwanted and not of biological interest, while we necessarily modify our notion of replicates, for the gene expression levels in single cells cannot be measured in replicate. To ensure that the right kind and amount of variation is removed from gene counts we estimate the effect of unwanted variation on these counts using suitably defined *pseudo-replicates* of cells or *pseudo-cells* that have the same biology, and we propose strategies to define pseudo-replicates. Using five publicly available datasets, we compare RUV-III-NB to several popular methods for normalizing scRNA-seq data and demonstrate its ability to retain biological signals and remove unwanted variation both in terms of *cell embedding* and gene-level count data, when biology and unwanted variation are associated.

## Results

### RUV-III-NB preserves biology when library size and biology are associated

In NSCLC study, the library size is associated with biology because the large epithelial cells have larger library sizes than those of the immune cells, and among the immune cells, monocytes are the largest, and they also have the largest average library size (Fig. 1A). RUV-III-NB identified log library size as a source of unwanted variation (Supp. Fig. 1A) and managed to separate the larger monocytes from the rest of the immune cells (Fig. 1B) better than sctransform-log corrected data (Fig. 1A) and other methods (Supp. Fig. 2). The silhouette statistic (Fig. 1C) shows that RUV-III-NB log PAC and Dino are the only normalization methods that improve the biological signals over that of the simple scran normalization. Apart from enhancing biological signals, a good normalization method should reduce effects of the unwanted factors in the normalized data. To investigate this, within cells of the same type, we examine the remaining effects of the library size in the normalized data using several metrics. Figure 1D shows that the leading principal components of sctransform-Pearson, RUV-III-NB log PAC and Dino-normalized data have the least association with log library size, with RUV-III-NB normalized data consistently having the lowest correlation with log library size across all genes (Supp. Fig. 3A). RUV-III-NB log PAC also produces median and IQR of relative log expression (RLE) that have the least association with log library size (Fig. 1E) and the smallest proportion of differentially-expressed genes (DEG) when cells with below and above median log library size are compared (Fig. 1F). Looking across all metrics, RUV-III-NB clearly has the best overall performance. Not only does it enhance the biological signals, it is also the most successful in removing library size effects from the data and in the differential expression analysis between cells of differing library sizes (Fig. 2).

**Fig. 1:**
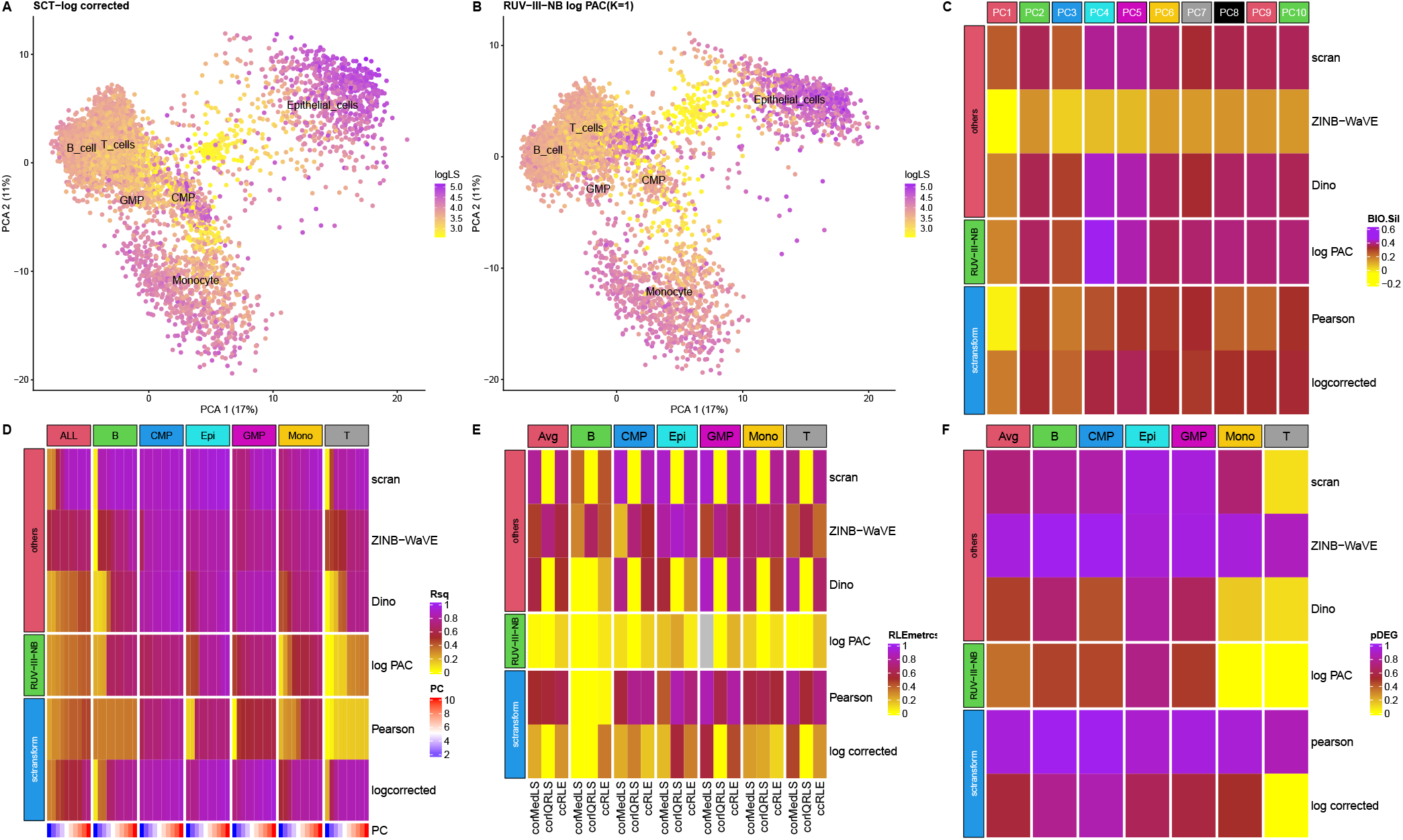
NSCLC study. (A) PC of sctransform-log corrected count. Colour refers to log library size. (B) PC of RUV-III-NB log percentile adjusted count (PAC). These show that monocytes are better separated from the rest of the cells. (C) Biological silhouette. RUV-III-NB and Dino are the only methods that improve the biological silhouette over that for scran normalization. (D) Heatmap of R-squared between logLS and PC of normalized data. RUV-III-NB and sctransform-Pearson have the lowest correlation, with RUV-III-NB still retaining some of the size-related heterogeneity within a cell type. (E) Correlation and canonical correlation (ccRLE) between median and IQR of relative log expression (RLE) and log library size. (F) Proportion of DEG between cells with below and above median log library size.

**Fig. 2:**
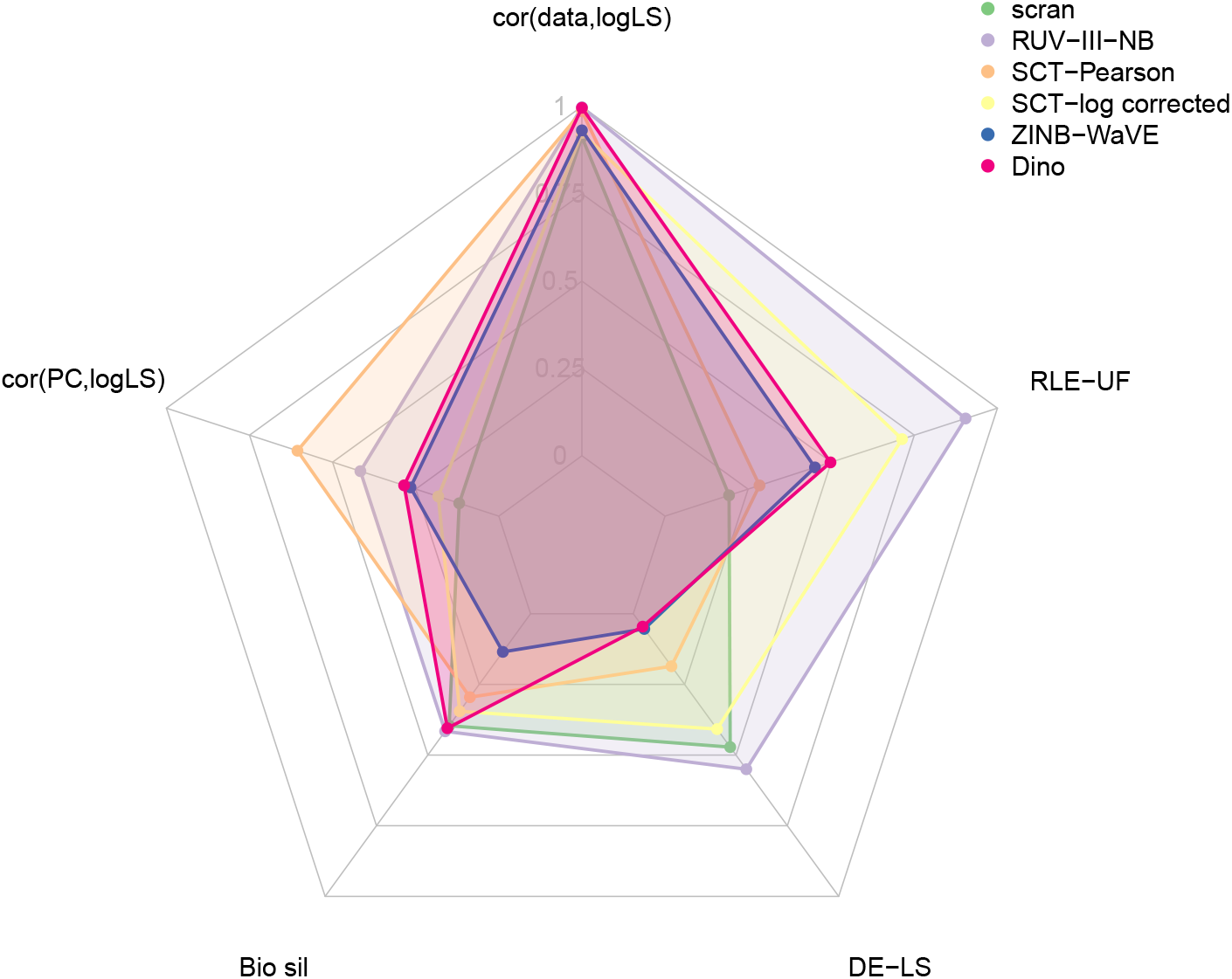
Overall performance of normalization methods in the NSCLC study. Each vertex corresponds to a metric and the length of the shaded area corresponds to level of performance with respect to the metric. For assessment metrics where lower indicates better performance such as technical (batch) silhouette and correlation between RLE characteristics and unwanted factors, the length of the segment is calculated as 1-metric. All metrics, except for biological silhouette, are calculated as an average of within cell-type statistics.

### RUV-III-NB preserves biology when batch and biology are associated

In the cell line study, there are two cell types but the cell types were sequenced in different pairs of the three batches. This creates an association between biology and batch. RUV-III-NB identified log library size (Supp Figs. 1 B) and batch (Supp Figs. 1 C) as major sources of unwanted variation. After scran normalization, the leading PC still clearly exhibit library size (Fig. 3A) and batch effects (Fig. 3B). RUV-III-NB removes the batch effects from the leading PC (Fig 3C) as does scMerge (Supp. Fig. 4). MN-NCorrect, Seurat3-Pearson, Seurat3-log corrected and ZINB-WaVE do not remove the batch effects, while fastMNN and Seurat3-Integrated remove the batch effects but also remove biology (Supp. Fig. 4). Only RUV-III-NB and scMerge improve the biological signals when compared with simple scran normalization (Fig. 3D), with RUV-III-NB being slightly better at reducing correlation between the normalized data and log library size (Supp. Fig. 3B), and much better at removing the effect of the unwanted factors from the RLE (Fig. 3E) and from the differential expression analysis Fig. 3D). Considering all the different metrics together, we see a clear advantage of RUV-III-NB and scMerge over the other methods, and an advantage of RUV-III-NB over scMerge for the RLE and differential expression metrics (Fig. 4).

**Fig. 3:**
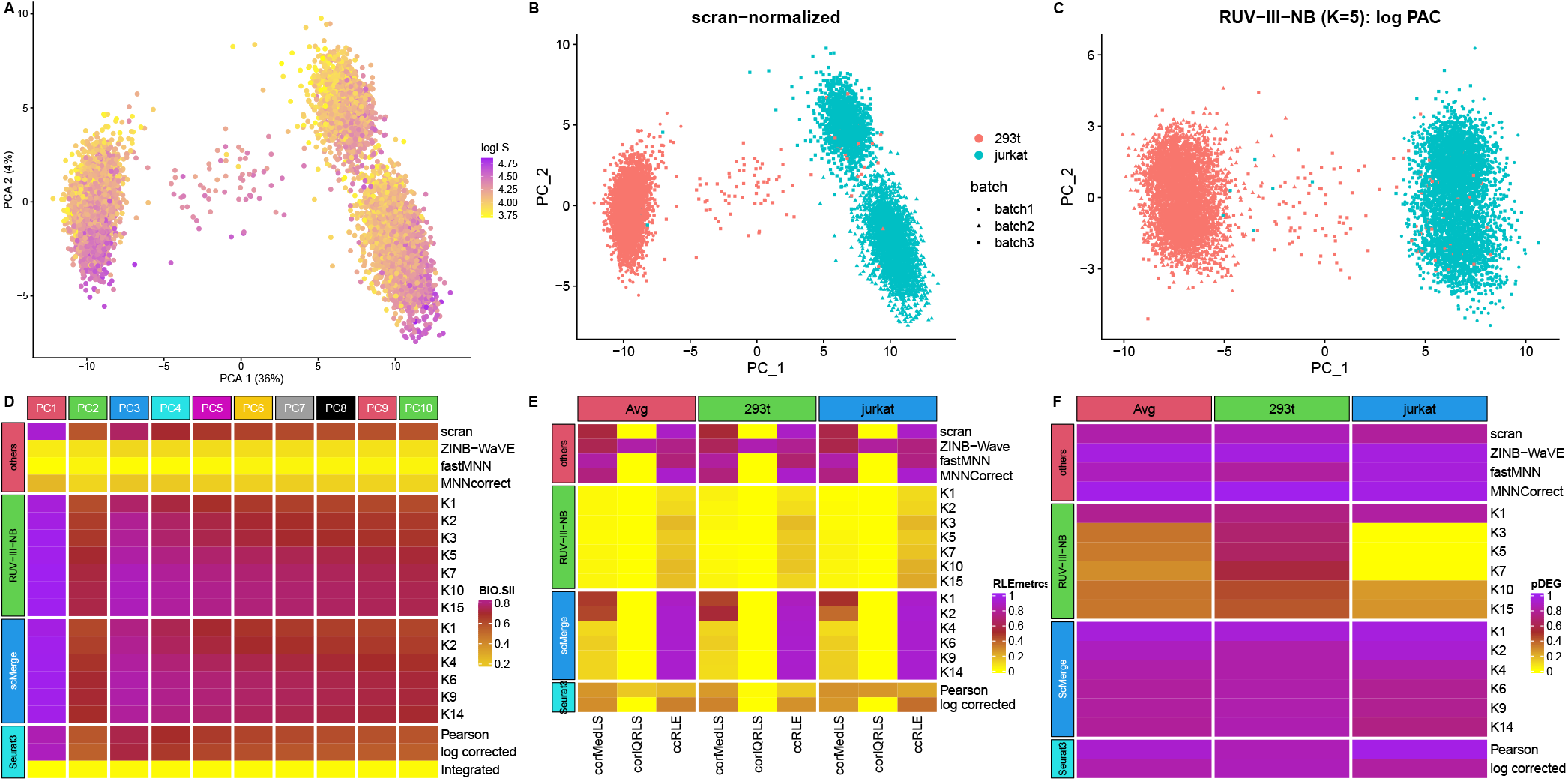
Cell line study. (A) The first two PC of scran-normalized data. Colour refers to log library size. The library size effect is clearly visible in the PC. (B) PC of scran-normalized data. Colour refers to cell type. Batch effects are visible for the Jurkat cells. (C) PC of RUV-III-NB log percentile adjusted counts (PAC). Clustering by cell type is clearly visible with batch effects removed. (D) Average biological silhouette score. RUV-III-NB and scMerge improve the biological signal and increasing the number of unwanted factors beyond a certain point only slightly degrades performance. (E) Correlation between median and IQR of relative log expression (RLE) with log library size and canonical correlation (ccRLE) between median and IQR of RLE and log library size and batch. (F) Proportion of DEG when comparing cells of the same type across batches.

**Fig. 4:**
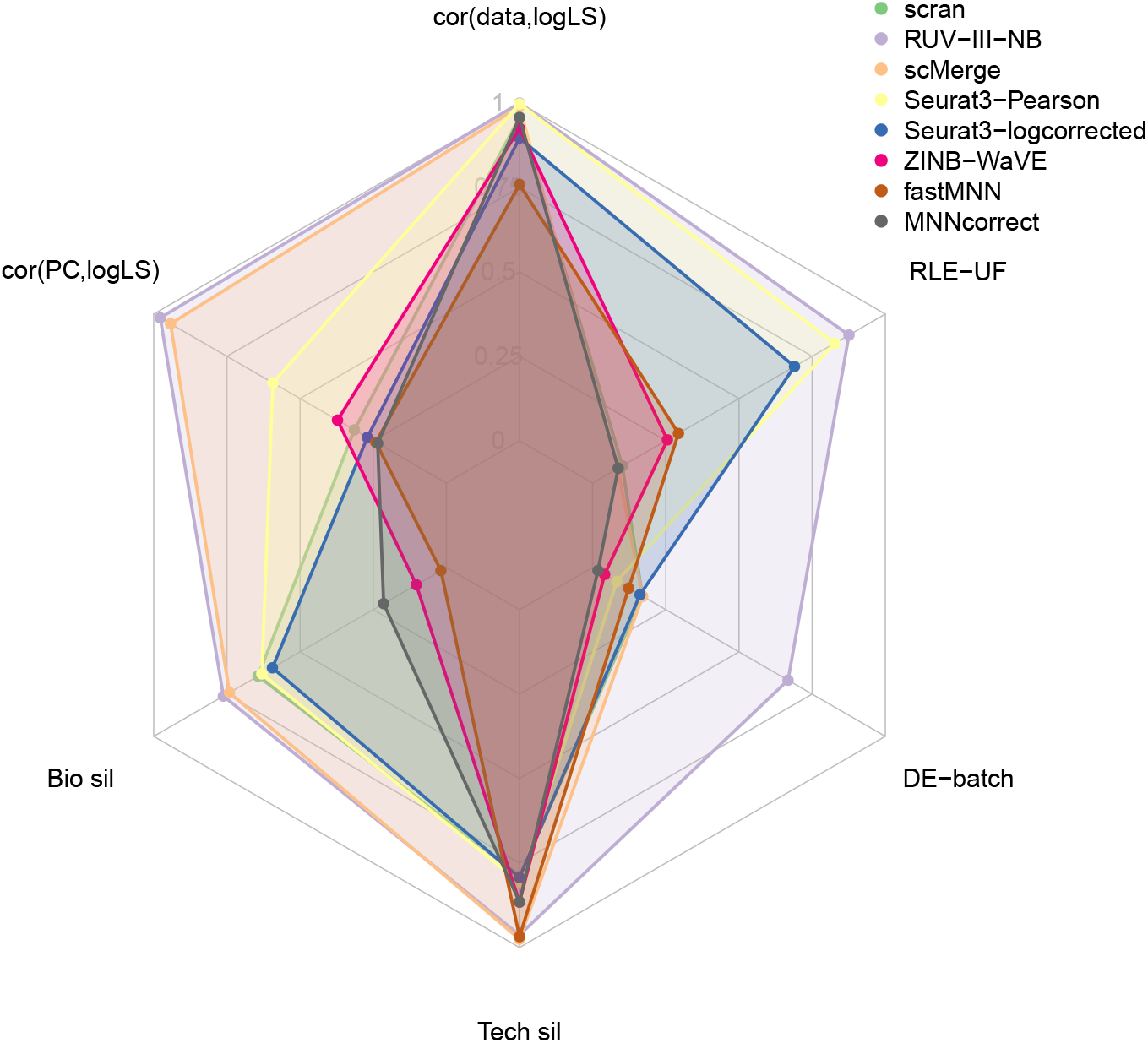
Overall performance of normalization methods in the celline study. Each vertex corresponds to a metric and the length of the shaded area corresponds to level of performance with respect to the metric. For assessment metrics where lower indicates better performance such as technical (batch) silhouette and correlation between RLE characteristics and unwanted factors, the length of the segment is calculated as 1-metric. All metrics, except for biological silhouette, are calculated as an average of within cell-type statistics.

The ability of RUV-III-NB to preserve biological signals and its excellent performance in terms of the RLE and differential expression metrics is also observed in the CLL study (Supp. Figs. 5, 7, 12B, 15), another study with UMI count where biology and batch are associated. However, in the Gaublomme study that does not have UMI counts, scMerge is slightly better than RUV-III-NB for almost all metrics, including the RLE and differential expression analyses (Supp. Figs. 6, 8 and 12C).

### RUV-III-NB preserves biology when biology is not associated with unwanted factors

The statistical model behind RUV-III-NB is designed so that removal of unwanted variation takes into account their potential association with biology. It is therefore of interest to examine how RUV-III-NB fares when the unwanted factors and biology are not associated. In the pancreas study, the eight cell types are present in both of the batches that correspond to different technological platform, and within each platform there is little difference in the average library size distribution between cell types (Supp. Fig. 10). Thus, there is only small amount of association between unwanted factors, in this case log library size and batch (Supp. Figs. 1H and I), with biology.

The leading PC of the scran-normalized data shows that cells of the same type are split by their batch of origin (Supp. Fig 11A). RUV-III-NB, scMerge and Seurat3-Integrated integrate the two batches well so that cells of the same type are clustered together (Supp. Figs. 11D,F and I). RUV-III-NB, together with scMerge and Seurat3-Pearson consistently manage to reduce the correlation between normalized data and log library size for homogeneous cell types (Fig. 5A). Seurat3-Integrated, RUV-III-NB and scMerge are the most successful in improving biological signals (Fig. 5B). But in terms of *R*^2^ between leading PC and log library size (Fig. 5C) and technical silhouette (Fig. 5D), scMerge and Seurat3-Integrated are slightly better than RUV-III-NB. This suggests that the more cautious approach of RUV-III-NB slightly reduces its ability to remove unwanted factors from the embedding, although RUV-III-NB is still the best method for removing the effect of unwanted factors from the normalized data, resulting in better RLE and differential expression analysis (Figs. 5E and F). When all metrics are considered together, RUV-III-NB still has the best overall performance (Fig. 6).

**Fig. 5:**
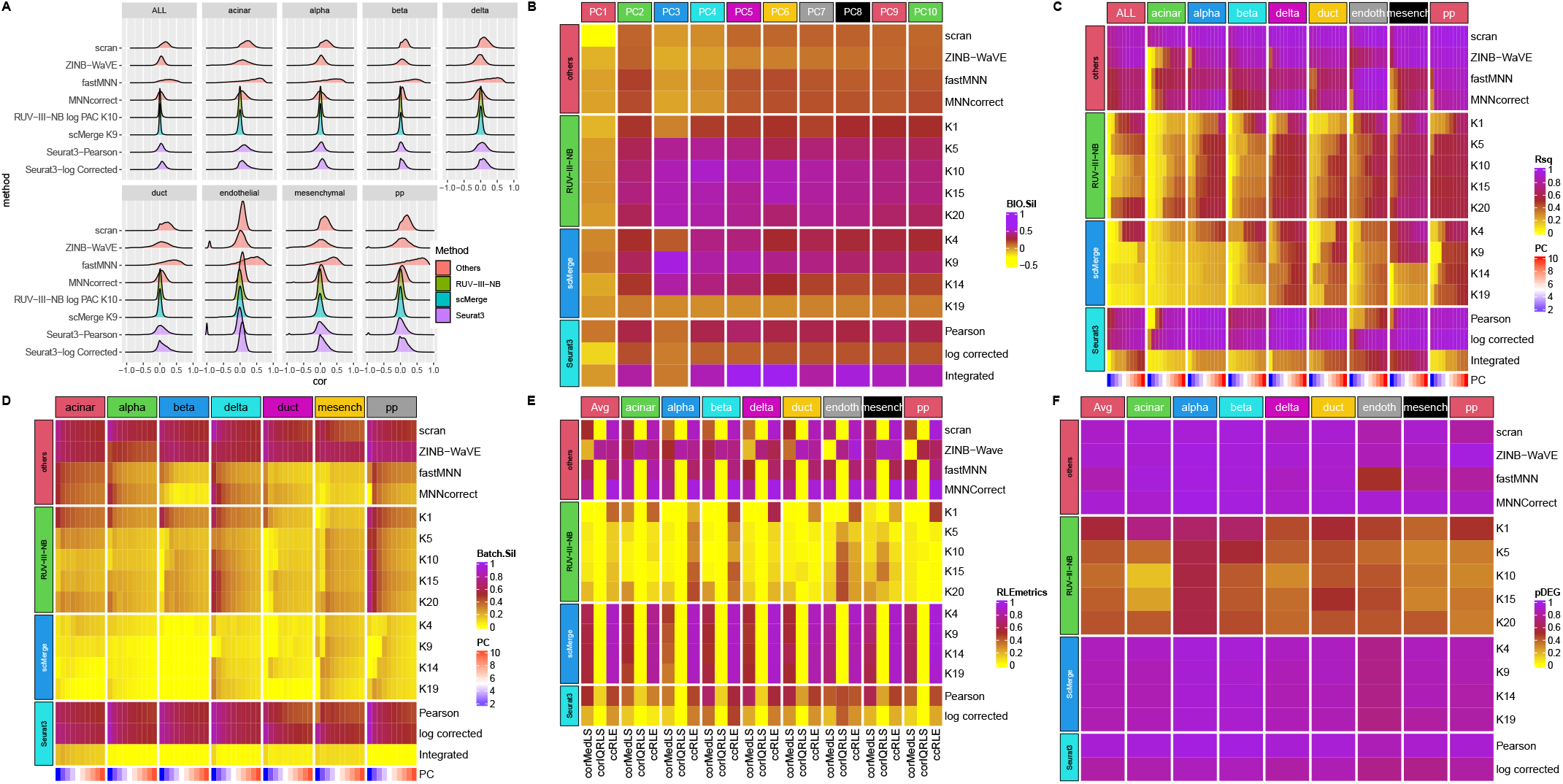
Pancreas study. (A) Densities of Spearman correlations between log library size and normalized data for ALL and each cell type. RUV-III-NB has the most concentrated density around zero, followed by scMerge. (B) Biological silhouette score. Seurat3-Integrated has the best biological silhouette score, followed by RUV-III-NB. (C) Heatmap of R-squared between logLS and PCs of normalized data. RUV-III-NB and scMerge have the lowest correlation, with RUV-III-NB still retaining some of the size-related heterogeneity within a cell type. (D) Technical silhouette scores for each cell-type. scMerge has the lowest silhouette, followed by RUV-III-NB. (E) Correlation between median and IQR of relative log expression (RLE) with log library size and canonical correlation (ccRLE) between median and IQR of RLE and log library size and batch. (F) Proportion of DEG when comparing cells of the same type across batches.

**Fig. 6:**
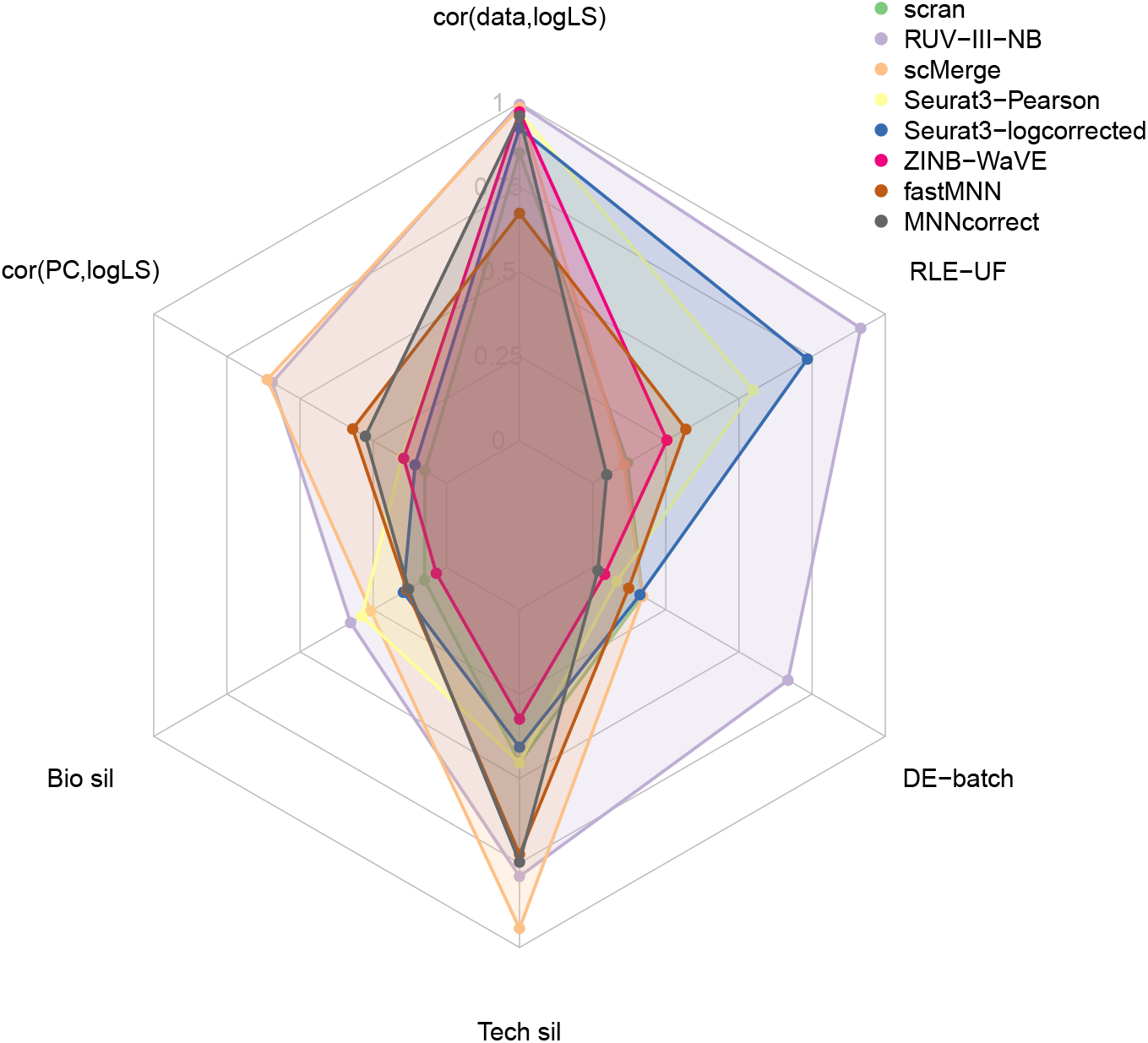
Overall performance of normalization methods in the Pancreas study. Each vertex corresponds to a metric and the length of the shaded area corresponds to level of performance with respect to the metric. For assessment metrics where lower indicates better performance such as technical (batch) silhouette and correlation between RLE characteristics and unwanted factors, the length of the segment is calculated as 1-metric. All metrics, except for biological silhouette, are calculated as an average of within cell-type statistics.

### RUV-III-NB accommodates size heterogeneity within a cell type

With UMI counts the library size corresponds closely to the number of molecules inside a cell and hence cell size. If there is size-related heterogeneity among cells of the same type, library size in experiments with UMI is biologically meaningful. We investigate the ability of the different normalization methods to isolate these biologically meaningful library size effects from the unwanted (technical) library size effects. To do this, for the NSCLC study we performed DE analysis comparing monocytes with smaller (< median) vs larger (≥ median) library size. The results show that RUV-III-NB has the lowest proportion of DEG (Fig. 1F), which suggests that RUV-III-NB removed the unwanted library size effects most effectively. We then performed KEGG pathway analysis among the DEG to investigate whether the DEG obtained are biologically meaningful. We found that only DEG from RUV-III-NB log PAC and sctransform-log corrected were significantly enriched with terms from the phagosome pathway (Supp. Fig. 13). This is consistent with^15^ who reported that larger monocytes have increased phagocytic activity. We carried out a similar analysis for the pancreas study where we compared beta cells with above and below median library sizes from the inDrop experiment.^16,17^ reported that patients with type II diabetes have reduced beta cells size. We found that that only the DEG from RUV-III-NB log PAC were significantly enriched with terms from the insulin resistance pathway (Supp. Fig. 14). We conclude that only RUV-III-NB normalization can reliably reveal size-related heterogeneity among cells of the same type.

### RUV-III-NB improves concordance with ‘gold standard’ DEG

For the Cell line, Gaublomme and Pancreas studies, we also compared the concordance of DEG based on data normalized by the different methods with the ‘gold standard’ DEG. For the Cell line study, the DEG are from the 293T vs Jurkat cell comparison, for the Gaublomme study we compare pathogenic vs sorted non-pathogenic Th-17 cells, while for the Pancreas study we compare alpha and beta cells. We found that for the Cell line and Gaublomme studies where batch is associated with biology, RUV-III-NB has the best concordance (Figs. 7A-B), while for the Pancreas study (Fig. 7C) where batch and biology are not associated, none of the batch-effect removal methods improve on scran normalization, with RUV-III-NB ranking second after Seurat3 with log-corrected counts.

**Fig. 7:**
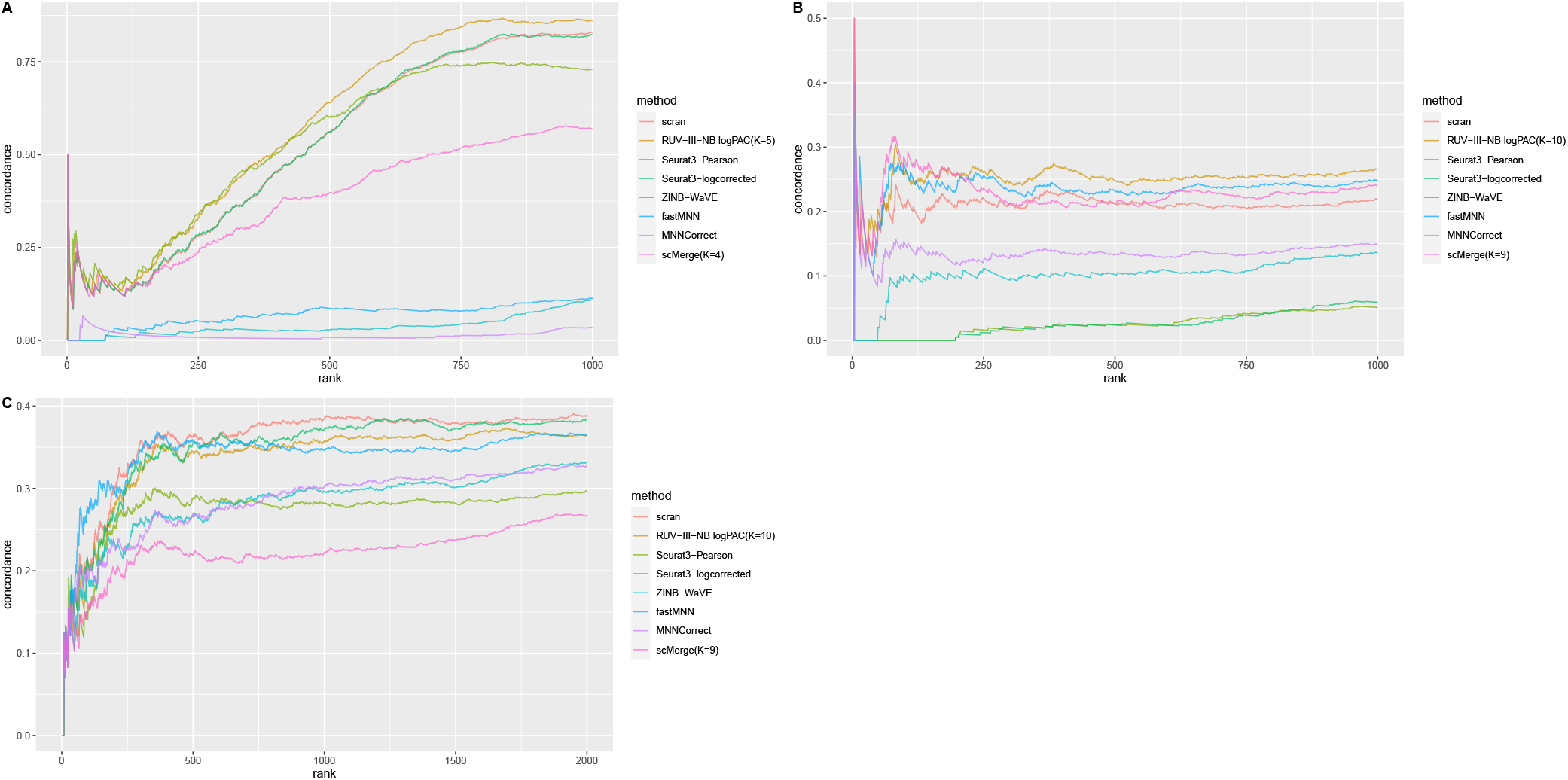
Concordance of DEG. (A) Jurkat cells in the cell line study. RUV-III-NB has the best concordance, followed by Seurat3. (B) Pathogenic vs Sorted Non-Pathogenic cells in the Gaublomme study. RUV-III-NB has the best concordance followed by fastMNN and scMerge. (C) Alpha vs Beta cells in the Pancreas study. scran has the best concordance followed by Seurat3 and RUV-III-NB.

### RUV-III-NB performance is robust

The RUV-III-NB algorithm require users to specify the negative control gene set and the number of unwanted factors. Using the cell line dataset, we investigate the sensitivity of the key performance metrics against these parameters. We use five different strategies to identify the negative control gene set and varying *K* from 1 to 20. Supp. Fig. 16A demonstrate that for four negative control gene sets, including set 2 that uses the default single-cell housekeeping genes, the R^2^ between log library size and leading principal component of normalized data is relatively robust when *K* is increased and thus potentially overestimated. Set 4, in which the negative control gene set was identified as non-DEG from the batch with two cell lines (batch 3), is the only one where the R^2^ is affected by overestimation of *K*. In terms of average batch (Supp Fig. 16B) and biological silhouette width (Supp Fig. 16C), its performance is quite similar across different negative control gene sets, for *K* ≥ 2.

### Computing time

The original implementation of RUV-III-NB requires a High-Performance Computing (HPC) environment. For the examples used in this paper, the running time on an HPC environment with 15 cores and 120 Gb total RAM (8Gb RAM per core), ranges from approximately 120 minutes for the CLL dataset with around 1,650 cells to around 280 minutes for the Pancreas dataset with more than 10,000 cells (Supp. Fig. 17A). The running time is approximately a square root, rather than a linear function of the number of cells. Studies involving scRNA-seq are growing in size and it is now not uncommon to have studies with several hundred thousands of cells. To meet this challenge, we also provide a fast implementation of RUV-III-NB, which we call *fastRUV-III-NB*. For *K* ≤ 10, the fast implementation is faster than MNNCorrect and scMerge and about half as fast as Seurat3 (Supp. Fig. 17B). Importantly, judging from several key metrics (Supp.Fig.18), *fastRUV-III-NB* achieves the same level performance as the original RUV-III-NB. The speed-up is achieved primarily by estimating gene-level parameters using a subset of cells (default = 20%). To reduce memory requirements *fastRUV-III-NB* processes the data as a `DelayedArray` object.

## Discussion

Single-cell RNA-seq offers us an unparalleled opportunity to advance our understanding of the transcriptome at the single cell level. However, scRNA-seq data contains significant amounts of unwanted variation that, when left unaddressed, may compromise downstream analyses. Most methods for removing unwanted variation from scRNA-seq data implicitly assume that the unwanted factors are at worst weakly associated with the biological signals of interest. In this paper, we have proposed RUV-III-NB, a statistical method for normalizing scRNA-seq data which does not make this assumption. The method adjusts for unwanted variation using pseudo-replicate sets, which should ensure that it does not remove too much biology when biology and unwanted variation are associated. Using publicly available data from five studies we show this to be the case.

We have benchmarked RUV-III-NB against methods that return gene-level normalized data as well as lower dimensional embedding. Both metrics are equally important in scRNA-seq experiments. While embedding is important and useful for clustering-based analysis to identify cells with similar biology, gene-level normalized data is used to identify markers genes to characterize the clusters. We have shown the distinct advantage of RUV-III-NB for UMI data in terms of embedding and normalized data when the unwanted variation is associated with biology. When biology is not associated with unwanted variation, RUV-III-NB has similar level of performance to Seurat3 and scMerge in terms of embedding and better in terms of normalized data.

A strong feature of RUV-III-NB is that it returns a sequencing count after adjusting for the unwanted variation. We call this the *percentile-invariant adjusted count* (*PAC*). These adjusted counts can be used as input to downstream analyses such as differential expression (DE), cell-type annotation and pseudotime analyses. In this paper, we have shown that when used for DE analysis, it delivers good control of false discoveries and improved power to detect ‘gold standard’ DE genes. In the vignette that accompanies the R package, we also demonstrated how the adjusted counts can be used to perform cell-type annotation.

RUV-III-NB can be used for both data with and without UMI, but its improvement relative to other methods is especially evident for UMI data. When using RUV-III-NB users need to specify the number of unwanted factors in the data (*K*) and the set of negative control genes. We have shown that RUV-III-NB performance is relatively robust to overestimation of *K* and the choice of negative control gene sets. While RUV-III-NB is developed primarily to remove within-study batch effects, it can also be used to integrate datasets from different studies where platform difference is a major source of unwanted variation. Using the Pancreas study, we have shown that the performance of RUV-III-NB for data integration purposes is quite competitive.

## Methods

We describe the RUV-III-NB model and algorithm here, with more details can be found in the Supplementary Methods. RUV-III-NB takes raw sequencing counts as input and models the counts *y_gc_* for genes *g* and cells *c*, as independent Negative Binomial (NB), *y_gc_* ~ *NB*(*μ_gc_*, *ψ_g_*) or Zero-Inflated Negative Binomial (ZINB) random variables, *g* = 1,…, *G*, *c* = 1,…, *N*. Here we will only discuss the NB model for UMI data and leave the ZINB model for read count data to the Supplementary Methods section. Without loss of generality, we further assume there are *m* groups among the *N* cells with the same underlying biology within and different underlying biology across groups. We will refer to these groups as pseudo-replicate sets, that is, sets of cells whose members will be regarded as replicates for the purposes of normalization. Let ***y***_*g*_ = (*y*_*g*1_, *y*_*g*2_,…, *y_gN_*)^*T*^ be the vector of counts for gene *g* and ***μ***_*g*_ be its vector of mean (i.e. expected value) parameters under the NB model. We use a generalized linear model with log link function to relate these mean parameters to the unobserved unwanted factor levels captured by the matrix **W** while the biology of interest will be embodied in the matrix **M**, these being related by

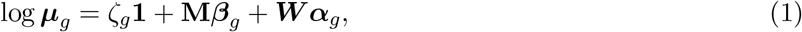

where ***M***(*N* × *m*) is the pseudo-replicate design matrix with ***M***(*c*, *j*) = 1 if the cth cell is part of the *j*th pseudo-replicate set and 0 otherwise, 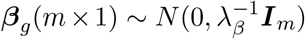 is the vector of biological parameters, with values for each of the m replicate sets, **W**(*N* × *k*) is the unobserved matrix of k-dimensional unwanted factor levels and 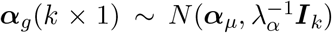 is the vector of regression coefficient associated with the unwanted factors, and finally *ζ_g_* is the location parameter for gene *g* after adjusting for unwanted factors, *g* = 1,… *G*. In our applications we found that setting λ_*α*_ = 0.01 and λ_*β*_ = 16 yield good results.

For a given number *k* of unwanted factors we use a double-loop iteratively re-weighted least squares (IRLS) algorithm, where in the inner loop, given current estimates of the dispersion parameters, we estimate the parameters of the loglinear model above, including the unobserved unwanted factor levels **W** (see Supplementary Methods for details). Once convergence is achieved there, we update the dispersion parameters in the outer loop. Two important constructs enable the algorithm to estimate the unobserved unwanted factor levels and their gene-specific effects on the sequencing count. These are the pseudo-replicate design matrix **M** and the set of negative control genes.

The pseudo-replicate design matrix **M** plays an important role for estimating the effect of the unwanted factors on the data.^14,18^ This effect is represented by ***α***_*g*_ and in RUV-III-NB it is estimated after projecting the current IRLS working vector onto the orthogonal complement of the subspace spanned by the columns of **M**. Given an estimate of ***α***_*g*_, we use the set of *negative control* genes to estimate the unobserved unwanted factor levels **W**. As stated above, negative controls are genes whose variation is (largely) unwanted and not of biological interest,^19^ i.e, ***β***_*g*_ ≈ 0 for all negative control genes *g*. The model for these genes thus reduces to

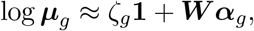

We recommend the use of single-cell housekeeping genes^20^ as the negative controls but users can (and may need to) devise their own negative control set. The important property of such genes is that they are affected by the same sources of unwanted variation as the other genes, and that their variation is not related to the biology of interest in the study.

### Strategies for defining pseudo-replicate sets

To estimate the effects of the unwanted variation on the gene counts, the RUV-III-NB algorithm requires users to specify one or more sets of cells with relatively homogeneous biology, and these are called *pseudo-replicate* sets. In cases where the biological factor of interest for each cell is known, e.g when different treatments are compared across the same cell type, or when two or more cell lines are being compared, then cells with the same level of the biological factor of interest can be declared to be a pseudo-replicate set. There will be situations where the biology of interest is not known a priori at the single cell level. For example, it is often the case that cell type information is unavailable in advance, especially for droplet-based technologies. For such situations we outline some strategies that can be used to define pseudo-replicate sets.

#### Single batch

When the data comes from a single batch, users can cluster the cells into distinct biologically homogeneous sets of cells. The clustering could be done using the log (normalized count + 1) where the scaling factor for normalization is calculated using `computeSumFactors` function in `scran` package.^1^ For clustering we recommend the use of a graph-based method such as the Louvain algorithm.^21^ Cells allocated to the same cluster can then be considered to form a pseudo-replicate set. We illustrate this strategy in Supp. Fig 19.

#### Multiple batches

When the data comes multiple batches, we need to match clusters containing cells with similar biology located in different batches. We recommend that users use the `scReplicate` function in the Bioconductor package `scMerge`^10^ for this purpose. This function takes log(normalized count + 1) as input and performs K-means clustering for each batch separately followed by identification of clusters in different batches that are mutual nearest clusters.^10^ Once these mutual nearest clusters (MNC) are identified, cells from the same MNC can be considered to form a pseudo-replicate set. We illustrate this strategy in Supp. Fig 20.

### Strengthening pseudo-replicate sets using pseudo-cells

Even when pseudo-replicate sets can be defined by clustering, the clustering may at times be imprecise, with considerable biological heterogeneity across cells in the same cluster. Thus declaring all such cells to be a pseudo-replicate set may risk removing some of the biological signal of interest. To address this issue, we introduce the idea of basing pseudo-replicate sets on *pseudo-cells*.

#### Pseudo-cells: single batch

Within a single batch and biology, we suppose that the major source of unwanted variation is library size, and that other intra-batch variation (e.g., well-to-well variation within a plate) is minimal. The idea is to form pseudo-replicates of pseudo-cells that have been constructed to have as much variation as possible in their library size while keeping their biology as homogeneous as possible, more homogeneous than we might see in actual single cells in a pseudo-replicate set. Suppose we have identified *m* pseudo-replicate sets using either known single cell biology or the strategy that we have just described above. For each of the pseudo-replicate sets, we form pseudo-cells that represent the pseudo-replicate set using the following pool-and-divide strategy:

1. Assign each cell to one of the *J* = 10 pools based on its library size, where pool *j* contains *n_j_* cells, *j* = 1… *J*.
2. Pooling: Let **Y**_*j*_ be the matrix of counts for cells belonging to pool *j* = 1, 2,… *J*, where rows corresponds to genes and columns corresponds to cells. We aggregate the counts for these cells by forming row totals of **Y**_*j*_ and denote the vector containing these row totals by **s**_*j*_ with components *s_gj_* = Σ*_c∈poolj_y_gc_*.
3. Dividing: For each gene *g*, we generate a count *z_gj_* using the pool-aggregated counts as follows: *z_gj_* | *s_gj_* ~ Binomial(*s_gj_*, *p* = 1/*n_j_*) where *s_gj_* is the aggregated count for gene *g* in pool *j* consisting of *n_j_* cells. This step is formally equivalent to randomly dividing the aggregated counts for the pool into counts for *n_j_* pseudo-cells and choosing one of the pseudo-cells at random. The hope is that the pseudo-cell so defined will exhibit average and so stabler biology in its gene counts, while concentrating the unwanted variation in the pool, here library size.
4. We thus obtain counts **z**_*j*_ = (*z*_1*j*_, *z*_2*j*_,…, *z_Gj_*)^*T*^ for the pseudo-cell that represents pool *j*.
5. We repeat steps 1-4 for all *J* pools and declare the *J* pseudo-cells so defined to be a pseudo-replicate set.
6. Finally, we carry out steps 1-5 above for the other pseudo-replicate sets, at the end of which we will have *m* pseudo-replicate sets each containing *J* pseudo-cells.

It can be shown that the counts assigned to these pseudo-cells will still have the quadratic mean-variance relationship typical of negative binomial random variables (see Supplementary Methods). The difference between these pseudo-cells and the real cells lies in the overdispersion parameter. For the same gene, the overdispersion parameter for pseudo-cells will be smaller, reflecting the reduced variability resulting from the pool-and-divide strategy. To incorporate this feature of pseudo-cells into the RUV-III-NB fitting process, we simply treat them as additional cells whose dispersion parameters are estimated separately from those of the real cells.

#### Pseudo-cells: multiple batches

When there are multiple batches, the procedure for forming pseudo-cells just described needs to follow the stratification of our cells into sets of MNC. Then we construct pseudo-cells for each of the clusters that makes up an MNC. For example, suppose we have *b* = 2 batches *A*, and *B* and we identified three clusters for each batch with the following MNC: (*A*_1_, *B*_2_), (*A*_2_, *B*_1_) and (*A*_3_, *B*_3_) where *A*_1_ refers to the first cluster in batch *A*, etc. The procedure for forming the pseudo-cells would then be as follows:

1. Start with the first MNC (*A*_1_, *B*_2_)
2. Assign each cell in *A*_1_ into one of the *J* groups based on its library size, where group *j* contains *n_j_* cells.
3. Pooling: Let **Y**_*j*_ be the matrix of counts for cells belonging to pool *j* where rows correspond to genes and columns corresponds to cells. Aggregate the gene counts in these cells by forming the row totals of **Y**_*j*_ and denote this new vector by *s_gj_*.
4. Dividing: For each gene *g*, we generate a count *Z_jg_* using the pool-aggregated counts as follows: *z_jg_* ~ Binomial(*s_jg_*, *p* = 1/*n_j_*) where *s_jg_* is the pool-aggregated count for gene *g*. As above, this step is equivalent to randomly dividing the aggregated counts for the pool into those for *n_j_* pseudo-cells and choosing one of the pseudo-cells randomly.
5. We thus obtain **z**_*j*_ = (*z*_1*j*_, *z*_2*j*_,…, *z_Gj_*) as the count data for pseudo-cell that represent pool *j*.
6. Repeat steps 2-5 for cells in *B*_2_.
7. Declare all the pseudo-cells formed in step 2-6 above to be a pseudo-replicate set.
8. Go to step 1 and repeat steps 2-6 for the second MNC (*A*_2_, *B*_1_) and third MNC (*A*_3_, *B*_3_)

When this procedure is completed, we will have as many pseudo-replicate sets as we have MNC sets and each pseudo-replicate set is made up of *b* × *J* pseudo-cells. We illustrate this strategy for *b* = 2 batches and *J* = 2 groups in Supp. Fig 21.

### Adjusted counts

Once we obtain the estimates of unwanted factors 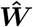 and their effects 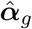, we remove their effects from the raw data. RUV-III-NB provides two forms of adjusted data. These adjusted data can be used as input to downstream analyses such as clustering, trajectory and differential expression analyses.

- Pearson residuals:

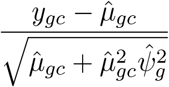

where 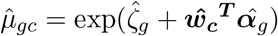. When *k* = 1 and 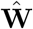 is approximately equal to log library size (up to a scaling factor), these Pearson residuals will roughly agree with those of,^4^ although different shrinkages of parameter estimates may lead to small differences. When *k* > 1 and some columns of **W** reflect batch effects, our Pearson residuals will also adjust for unwanted variation other than library size, such as batch effects.
- Log of percentile-invariant adjusted count (log PAC):

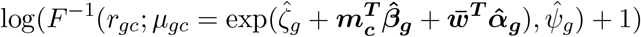

where *r_gc_* ~ *U*(*a_gc_*, *b_gc_*) and

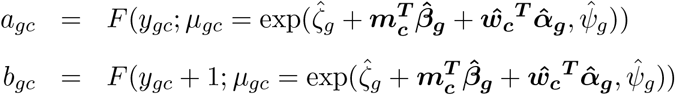

where *F*(·) is the negative binomial c.d.f and *F*^-1^(.) its inverse, ***m***_*c*_ is the *c^th^* row of the matrix ***M***, 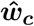 the *c^th^* row of the matrix 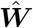 and 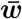 is vector of entries equal to the average level 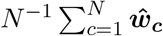 of unwanted variation. Here *U*(*a*, *b*) denoted a random variable uniformly distributed over the interval (*a*, *b*).

The intuition behind this adjustment is as follows. We first obtain the percentiles of the observed counts under the fitted NB model, where the mean value parameter includes terms for unwanted variation. Since negative binomials are discrete distributions, percentiles can only be determined up to an interval. To come up with an estimate of a percentile for practical use, we simply select a uniformly distributed random value from this interval in a manner suggested in.^22^ We then find the corresponding count for that estimated percentile under a different NB model, namely one where the mean parameter is free from unwanted variation, i.e. where 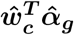 is replaced by 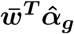. We then add 1 and *log*. Our definition of percentile-invariant adjusted count explicitly derives the counts as percentiles of a full NB distribution and in this regard it is similar to that in^23^ who proposed this approach to obtain batch-corrected bulk RNA-seq data. Their adjustment was only applied to non-zero counts, and left the zero counts intact. That was not expected to pose significant problems for bulk RNA-seq data where zero counts are relatively scarce, but because zero counts are very prominent in scRNA-seq data, we broaden their approach and also adjust zero counts. On the other hand, sctransform’s corrected count^4^ is calculated by taking away from the observed count the difference between the predicted counts at the observed and at the average log library size, followed by rounding to avoid non-integer values.

### Datasets for Benchmarking

To benchmark our methods against others, we use the following five datasets that encompass different technological platforms, illustrate different strategies for identifying pseudo-replicates and pose different challenges for normalization due to association between different unwanted factors and biology (Table 1).

**Table 1:**
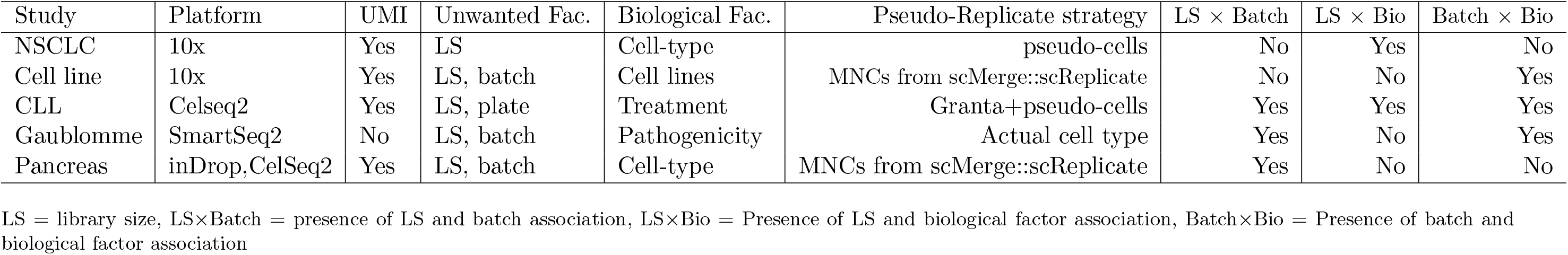
Characteristics of datasets used for benchmarking

Prior to normalization all datasets were subjected to quality control checks using Bioconductors’s `scater` package^24^ to remove low quality cells. Low abundance genes were also removed and additional parameters for each method were set to their default.

- Non-Small Cell Lung Cancer cells (NSCLC): The dataset was generated using 10x and is freely available from the 10x Genomics website (www.10xgenomics.com). The sequencing was done in one batch, so there should be no batch effects, but the cells are a mixture of cells with larger size such as epithelial cells and smaller cells such as T cells. The challenge here is to normalize when library size is associated with the biology, namely, cell-type. After QC, there were 10,0019 genes and 6,622 cells.
- Cell line: Here the 10x technology was used to sequence cells in three batches. One batch contained only the Jurkat cell line, another contained only the 293T cell line, while the third batch contained 50-50 mixture of both cell lines. Data were downloaded from 10x Genomics website. After QC, there were 7,943 genes and 9,027 cells.
- Chronic lymphocytic leukemia (CLL): This in-house dataset was generated using the CelSeq2 technology as part of a study investigating the transcriptomic signature of Venetoclax resistance. The cells were pre-sorted so that the vast majority are B-cells and were treated with dimethyl sulfoxide (DMSO) as well as single treatment (TRT) and combination treatments (TRT+) for one week, before being sequenced on six different plates. In addition to this, a small number of cells from the Granta cell line were included on each plate. After QC, there were 11,470 genes and 1,644 cells. The dataset is included as `CLLdata` object in the `ruvIIInb` R package.
- Gaublomme: Here Th17 cells derived under a non-pathogenic condition (TGF-*β*1+IL-6, unsorted: 130 cells from 2 batches and TGF-*β*1+IL-6; sorted for IL-17A/GFP+: 151 cells from 3 batches) and a pathogenic condition (Il-1*β*1+IL-6+IL-23, sorted for IL-17A/GFP+: 139 cells from 2 batches) were sequenced using the SMARTseq technology.^25^ After QC, there were 7,590 genes and 337 cells.
- Pancreas: Human pancreas islet cells from two different studies.^16^ used the inDrop technology to sequence the cells, while^26^ used the CELSeq2 technology. The datasets were downloaded from https://hemberg-lab.github.io/scRNA.seq.datasets/human/pancreas/. After QC, there were 11,542 genes and 10.687 cells.

### Benchmarking Methods

For the NSCLC study where there should be no batch effect and the only task is removing library size effects, we compared RUV-III-NB with the following methods: scran,^1^ sctransform,^4^ ZINBwave^12^ and Dino.^5^ For the other studies where batch effects are present, we compare RUV-III-NB to the following batch correction methods: mnnCorrect and fastMNN,^8^ Seurat3^11^ coupled with sctransform normalization, ZINBwave^12^ and scMerge.^10^ These methods have been selected because all of them return the gene-level normalized data required to calculate the benchmarking metrics (see below). This is in contrast with other methods such as Harmony,^9^ where the normalized data is only available as an embedding. Some of the methods produce multiple versions of normalized data and in Supp. Table 1, we provide details on which normalized data we used for calculating the various metrics in our benchmarking exercise.

We use the following criteria for assessing the performance of the different normalization methods:

- **Genewise correlations between the normalized data and log library size:** We expect a good normalization to remove any association between gene expression levels and log library size, especially when cells with the same biology are considered.
- **R**^2^ **between log library size and the leading PC:** For each cell-type, the coefficient of determination (R^2^) when regressing log library size on the leading PCs should be as low as possible. This is because we believe that a good normalization should reduce the association between the normalized data and log library size, so within a group of cells with similar biology, the leading PCs should contain little information about library size.
- **Silhouette statistics for clustering by batch (Technical Silhouette):** When batch effects are reduced, we should expect a lower degree of clustering by batch, within a set of cells with homogeneous biology.
- **Silhouette statistics for clustering by biology (Biological Silhouette):** When batch effects are removed, we expect biological signals to be strengthened and lead to better clustering by biology. For all methods, with the exception of Seurat3-Integrated, to calculate silhouette scores we used the first ten PC calculated using genes whose normalized expression variance lies in the top 50% and Euclidean distance. For Seurat3-Integrated, we use all anchor features for calculating PC. The number of anchor features is typically 2000, much less than the half of the total number of genes. When biological factors of interest are available from the dataset, these are used to calculate silhouette scores. Otherwise, we use the Bioconductor package `SingleR`^27^ to estimate the cell types. PC were derived using the R package `irlba`.^28^
- **Differential expression vs unwanted factors (DE-UF):** When comparing cells of the same cell-type across batches **(DE-batch)** or smaller vs larger library size **(DE-LS)**, a good normalization should *decrease* the proportion of differentially expressed genes (DEG).
- **Differential expression vs biology (DE-Bio):** When comparing cells across different biologies, a good normalization should increase the concordance between the results found with the current and those of an independent study, as measured by the number of DEG.
- **Canonical correlation between relative log expression medians and interquartile ranges (IQR) and log library size and batch variables (RLE-UF):** With good normalization, we expect that within a set of cells with homogeneous biology, the RLE plot summary statistics have little association with unwanted factors.Because the RLE calculation requires subtracting log of gene-specific median expression,^29^ the RLE plots were calculated using only genes with non-zero median expression.

### ‘Gold-standard’ DE genes

We compare the concordance of differentially-expressed genes (DEG) obtained from the different methods to the following ‘gold standard’ DEG:

1. Celline: ‘Gold standard’ DEG in this case were derived by comparing Jurkat and 293T cells from batch 3, which has cells from both cell lines. The assumption is that cells assayed in the same batch will exhibit similar batch effects that will, to some extent, cancel when we compare cells of different types. The DE analysis was performed using the Kruskal-Wallis test on the log(scran-normalized data + 1).
2. Gaublomme: ‘Gold standard’ DEG here were derived from an external dataset. We downloaded the raw Affymetrix CEL files from the GEO website (ID: GSE39820). The microarray data were normalized using the `GCRMA` package version 2.58.0 and DE analysis comparing non-pathogenic (TGF-*β* 1+IL-6) vs pathogenic (Il-1*β*1+IL-6+IL-23) microarray samples was performed using the `limma` package.^30^
3. Pancreas: ‘Gold standard’ DEG here were also derived from an external dataset. Normalized Agilent microarray expression data were downloaded from https://www.omicsdi.org/dataset/arrayexpress-repository/E-MTAB-465 and DE analysis comparing Alpha vs Beta cells was performed using `limma`.

## Data Availability

All datasets used in this paper are published datasets available for downloads from sources outlined in the Methods section above.

## Author Contributions

AS and TPS conceptualized the study, derived the model and the algorithm. AS and ADL implemented the algorithm. AS, JW and RM performed data analyses and interpretation. RT designed and generated data for the CLL study. AS wrote the manuscript with inputs from TPS, RM, JW, RT and ADL. All authors read and approved the submitted manuscript.

## Acknowledgement

The authors would like to thank Jean Yang, Yingxin Lin and Xiangnan Xu for their suggestions that have improved the quality of this manuscript.

