## Supplementary Figures and Tables for "RUV-III-NB: Normalization of single cell RNA-seq Data"

Agus Salim *et al.*

January 25, 2022

Supp. Table 1: Data Assays Used for Calculating Benchmarking Metrics

| Methods | Genewise Corr | Silhouette&PC | DE | RLE |
|---|---|---|---|---|
| fastMNN | reconstructed | reconstructed | reconstructed | reconstructed |
| MNNCorrect | corrected | corrected | corrected | corrected |
| scMerge | scMerge | scMerge | scMerge | scMerge |
| sctransfom-Pearson | y | y | y | y |
| sctransfom-log corrected | log(umi_corrected+1) | log(umi_corrected+1) | log(umi_corrected+1) | log(umi_corrected+1 |
| Seurat3-Pearson | scale.data@SCT | scale.data@SCT | scale.data@SCT | scale.data@SCT |
| Seurat3-log corrected | log(count@SCT+1) | log(count@SCT+1) | log(count@SCT+1) | log(count@SCT+1) |
| Seurat3-Integrated | NA | scale.data@Integrated | NA | NA |
| RUV-III-NB | logPAC | logPAC | logPAC | logPAC |
| ZINB-Wave | normalizedValues | normalizedValues | normalizedValues | normalizedValues |

* Consult the relevant R help files for each method to find what is contained in each assay.
Genewise Corr: Correlation between normalized gene expression and a factor e.g log library size, calculated gene-by-gene.
PC: Principal Component
DE: Differential Expression Analysis.
RLE = Relative Log of Normalized Expression.
PAC = Percentile adjusted count

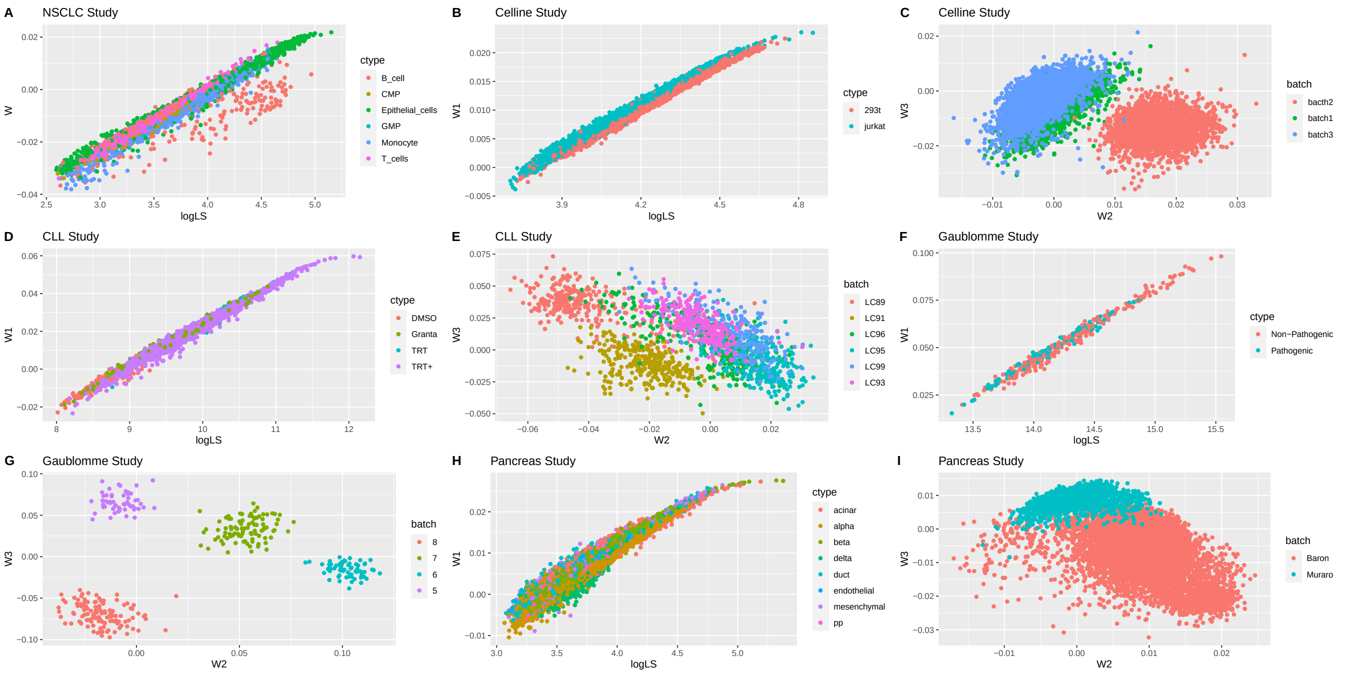


Supp. Fig. 1: Estimated unwanted factor values for the 5 studies

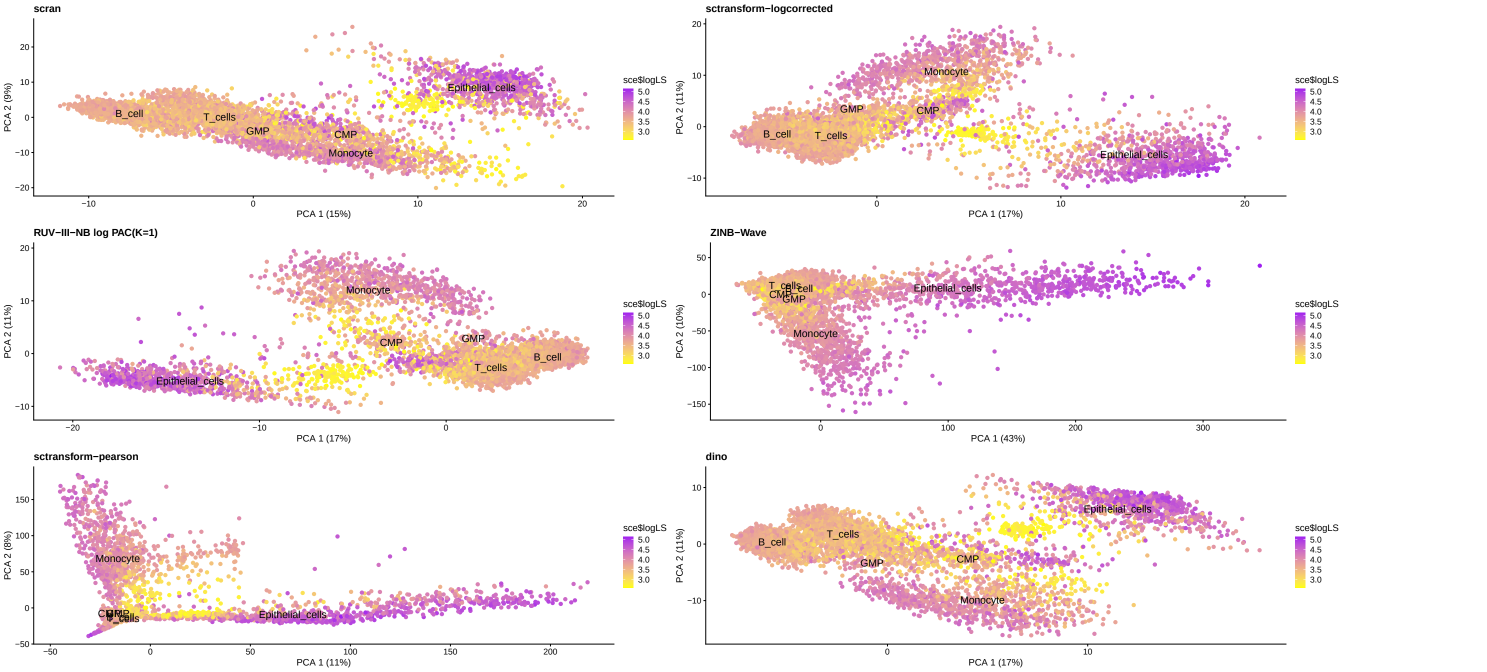


Supp. Fig. 2: NSCLC study. Leading PC of the differently normalized counts. Coloured by log library size.

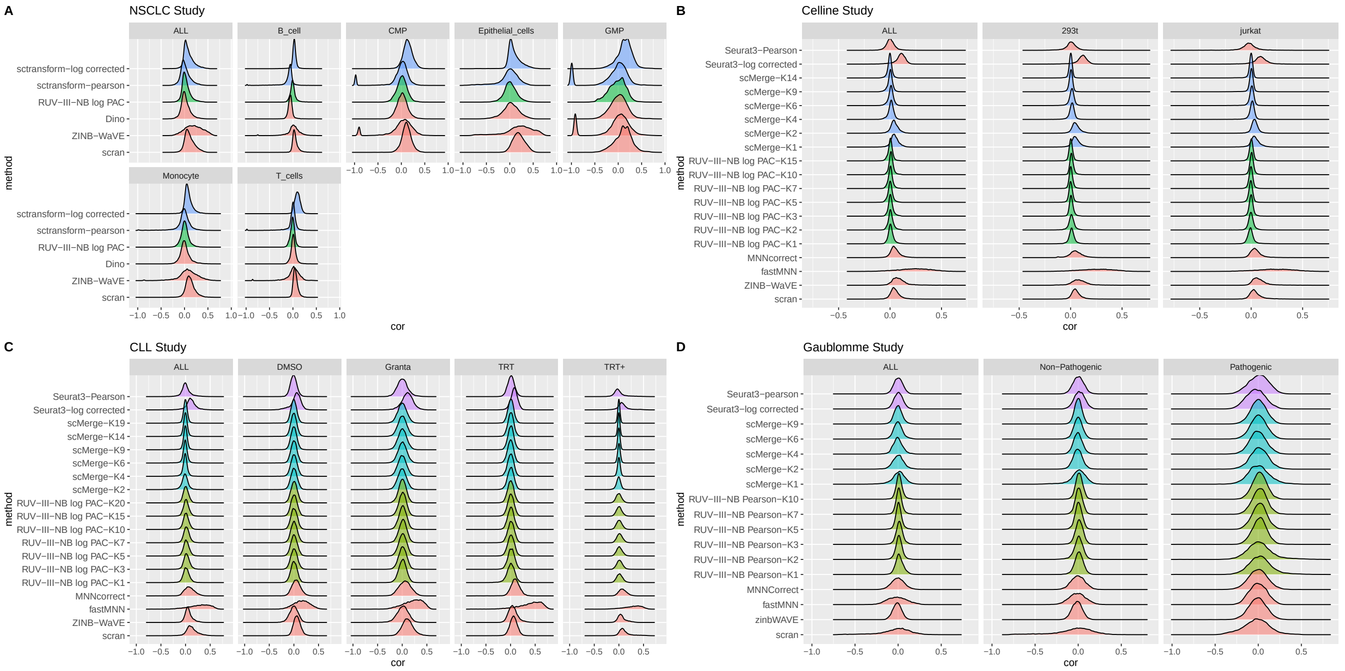


Supp. Fig. 3: Smoothed densities of gene count-log library size Spearman correlations for differently normalized counts. (A) NSCLC study. (B) Cell line study. (C) CLL study. (D) Gaublomme Study.

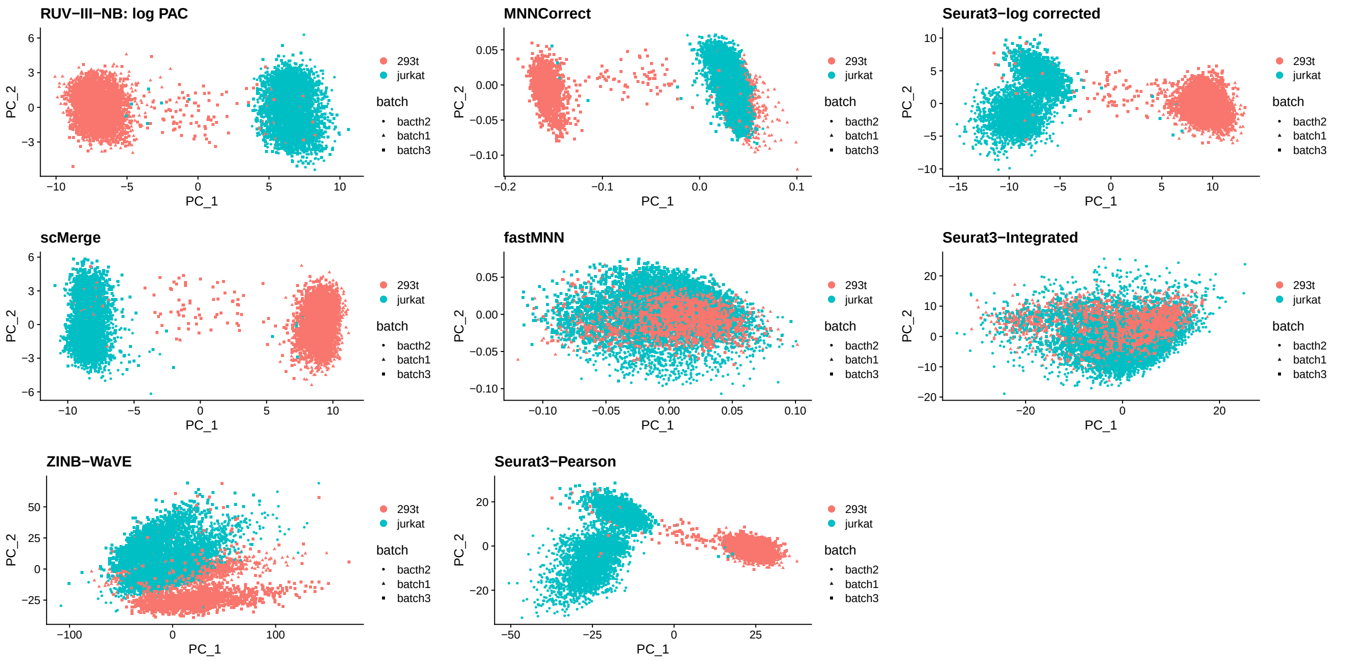


Supp. Fig. 4: Cell line study. Leading PC of the differently normalized data. Coloured by cell types.

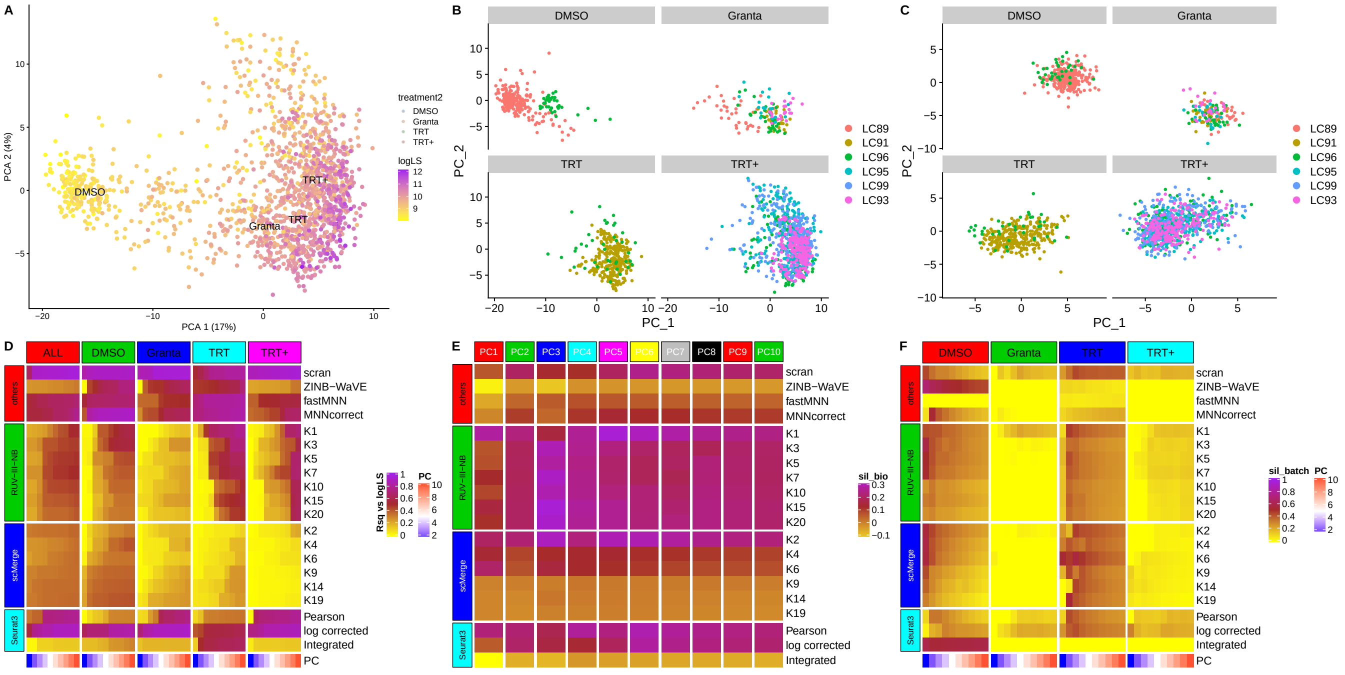


Supp. Fig. 5: CLL study.(A) The first two PC of scran-normalized data. Colour refers to log library size. The shallower sequencing of the DMSO-treated cells is clearly visible. (B) PC of scran-normalized data faceted by cell type. Colours refers to plate identity. The plate effects among DMSO and Granta cells are clearly visible. (C) PC of RUV-III-NB log PAC ($K = 20$) faceted by cell type. Colours refers to plate identity. No plate effects are visible. (D) Heatmap of R-squared between logLS and PC of normalized data. scMerge has the lowest correlation, followed by RUV-III-NB. (E) Biological silhouette scores. RUV-III-NB, scMerge and Seurat3 improve the biological silhouette score compared to scran, but scMerge's performance is sensitive to the choice of K. (F) Technical silhouette for each cell-type. RUV-III-NB and scMerge have the lowest silhouette score, followed closely by Seurat3.

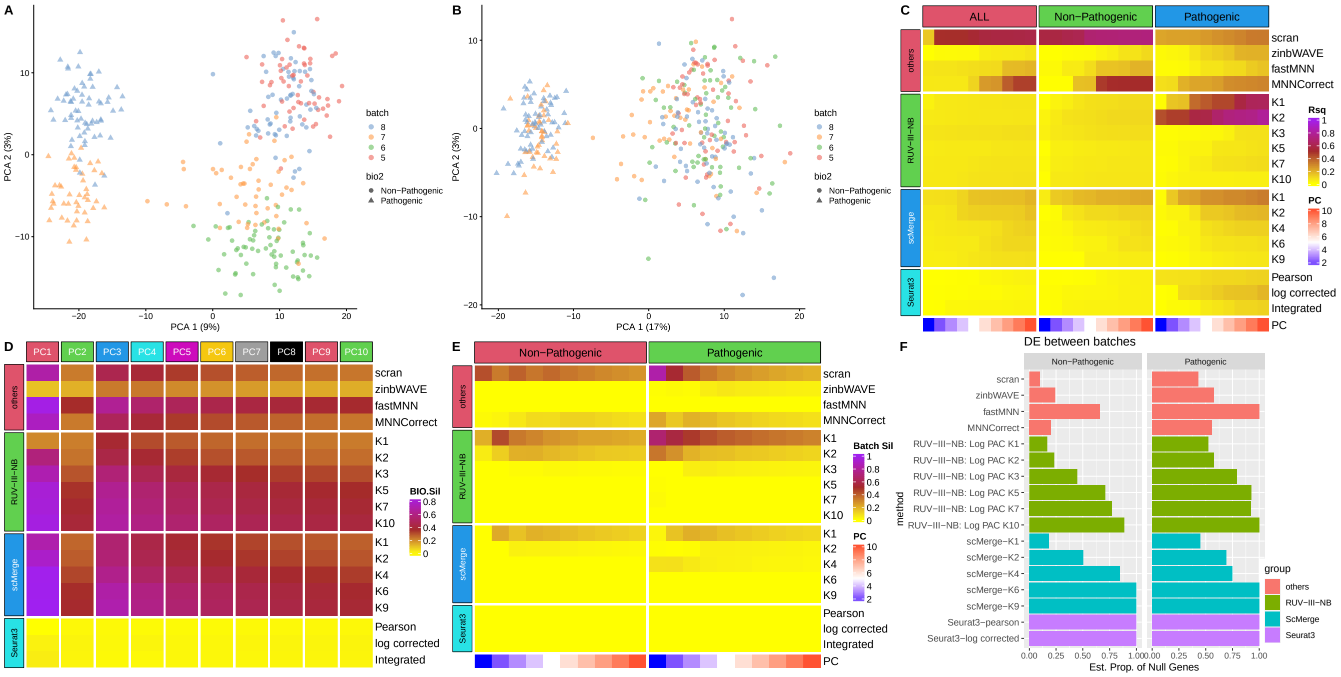

Supp. Fig. 6: Gaublomme study. (A) PC1 and PC2 of scran-normalized data. The first PC clearly shows the separation of pathogenic from non-pathogenic cells, but batch effects dominate the second PC. (B) PC of RUV-III-NB Pearson residuals ($K = 5$). Batch effect are no longer visible. (C) Heatmap of R-squared between logLS and PC of normalized data. All methods, except MNNCorrect reduce the correlation with logLS when compared to scran-normalized data. (D) Biological silhouette scores. RUV-III-NB improves the biological silhouette scores when compared to scran-normalization, followed by scMerge. The other methods reduce the biological signal in the process of removing unwanted variation. (E) Technical silhouette scores for each cell-type. All methods reduce the technical signal when compared with scran normalization. (F) Estimated proportion of non-DEG when comparing the same cell types across batches. Seurat3 and scMerge has the highest proportion of non DEG, followed by RUV-III-NB.

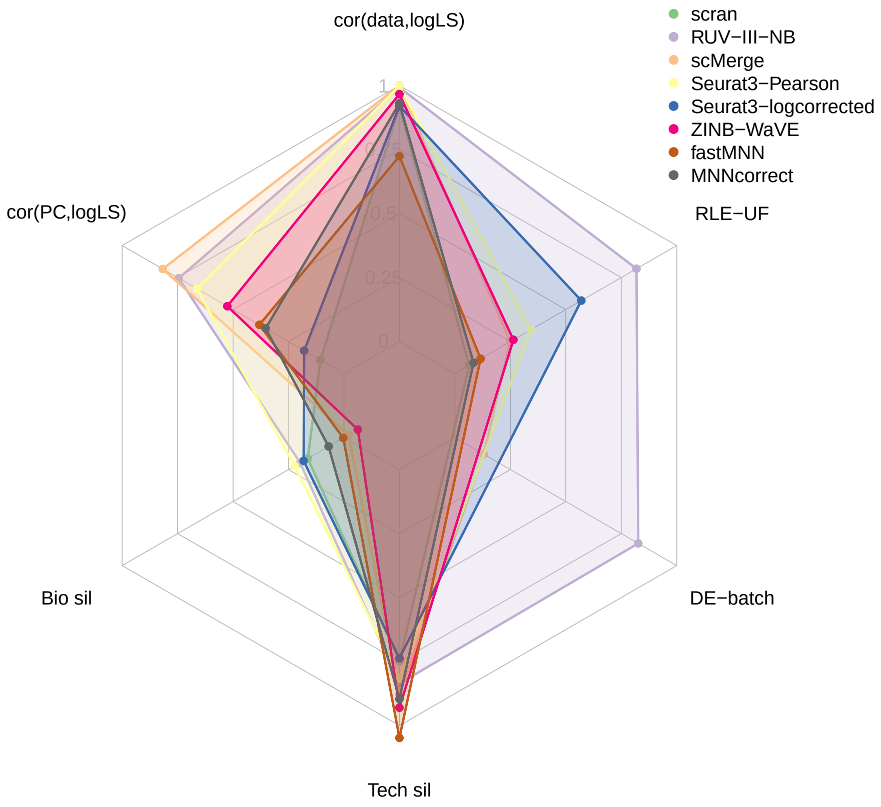


Supp. Fig. 7: Overall performance of normalization methods in the CLL study. Each vertex corresponds to a metric and the length of the shaded area corresponds to level of performance with respect to the metric. For assessment metrics where lower indicates better performance such as technical (batch) silhouette and correlation between RLE characteristics and unwanted factors, the length of the segment is calculated as 1-metric. All metrics, except for biological silhouette, are calculated as an average of within cell-type statistics.

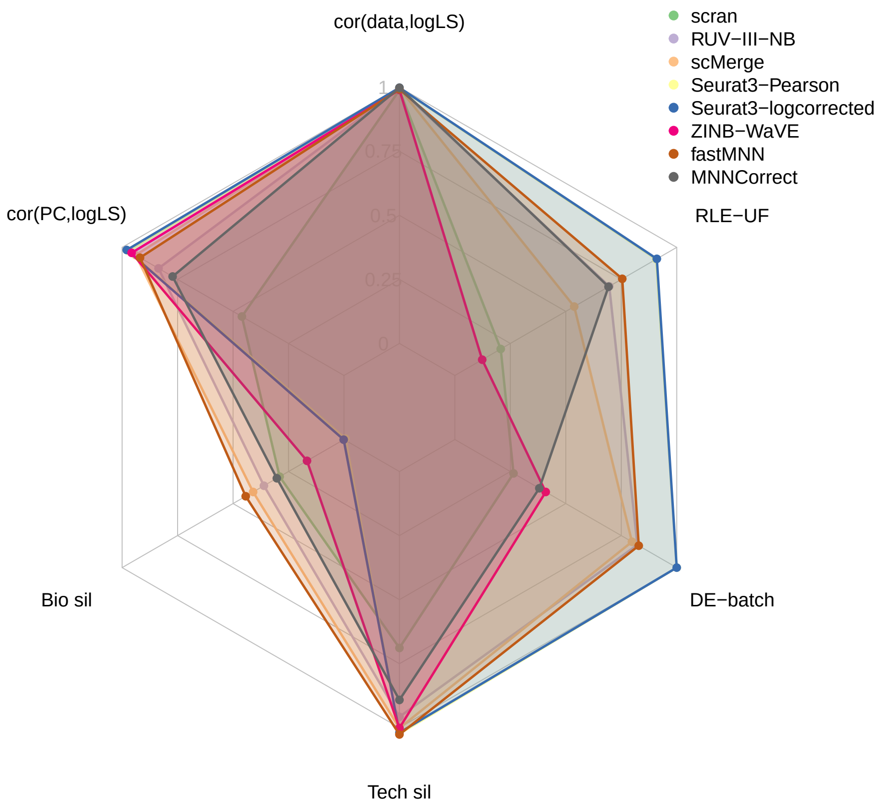


Supp. Fig. 8: Overall performance of normalization methods in the Gaublomme study. Each vertex corresponds to a metric and the length of the shaded area corresponds to level of performance with respect to the metric. For assessment metrics where lower indicates better performance such as technical silhouette and correlation between RLE characteristics and unwanted factors, the length of the segment is calculated as 1-metric. All metrics, except for biological silhouette, are calculated as an average of within cell-type statistics.

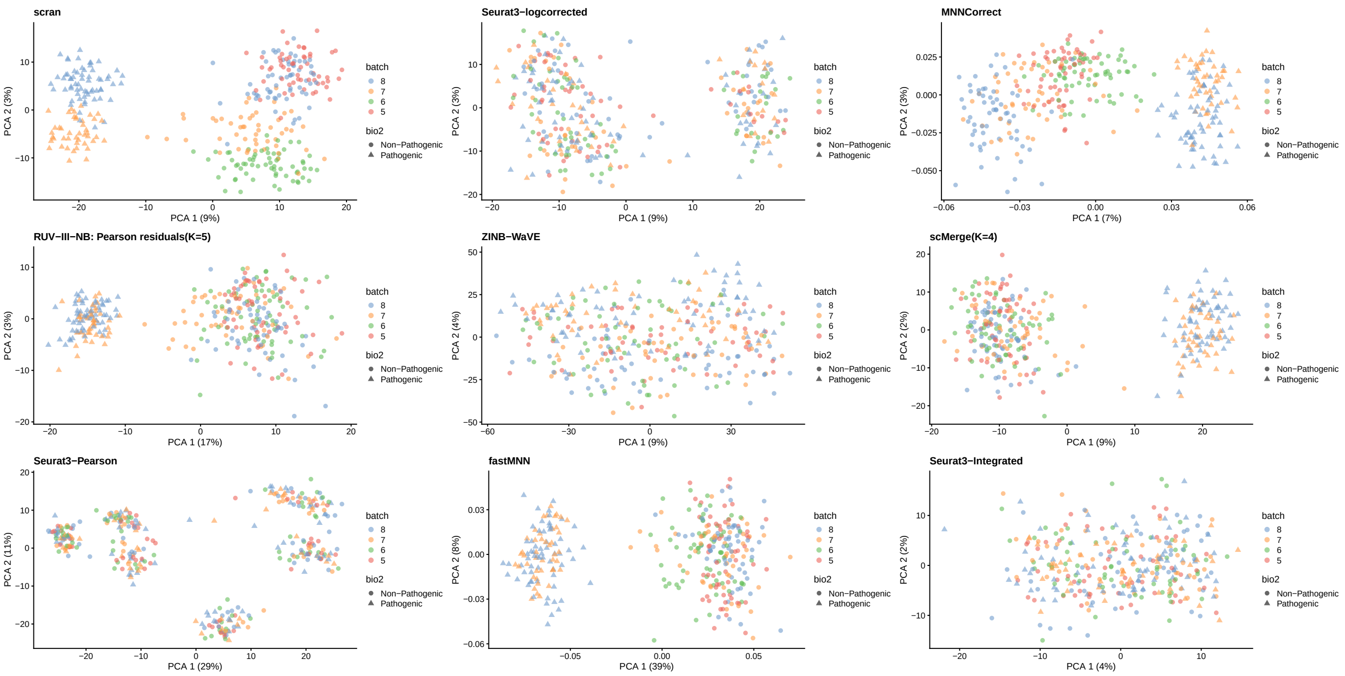


Supp. Fig. 9: Gaublomme study. Leading PC of the differently normalized counts. Coloured by batches, while shape refers to cell type.

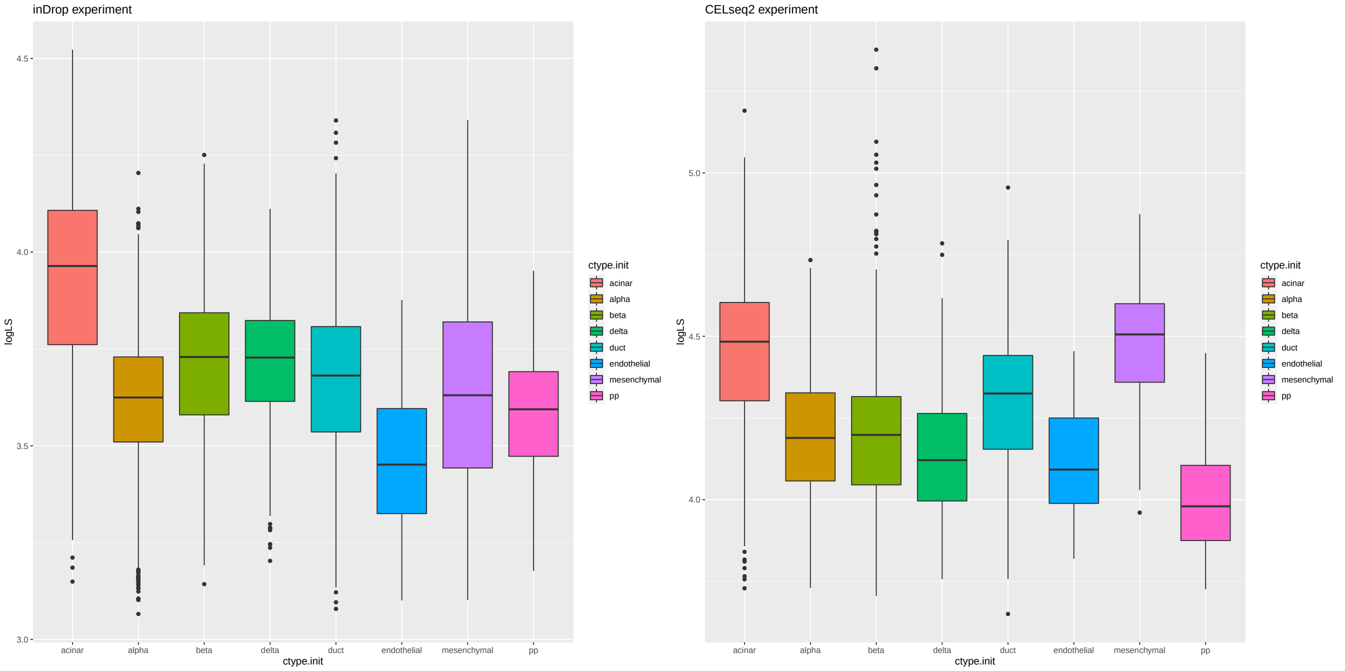


Supp. Fig. 10: Pancreas study. Log library size distribution for the different cell types in the two experiments.

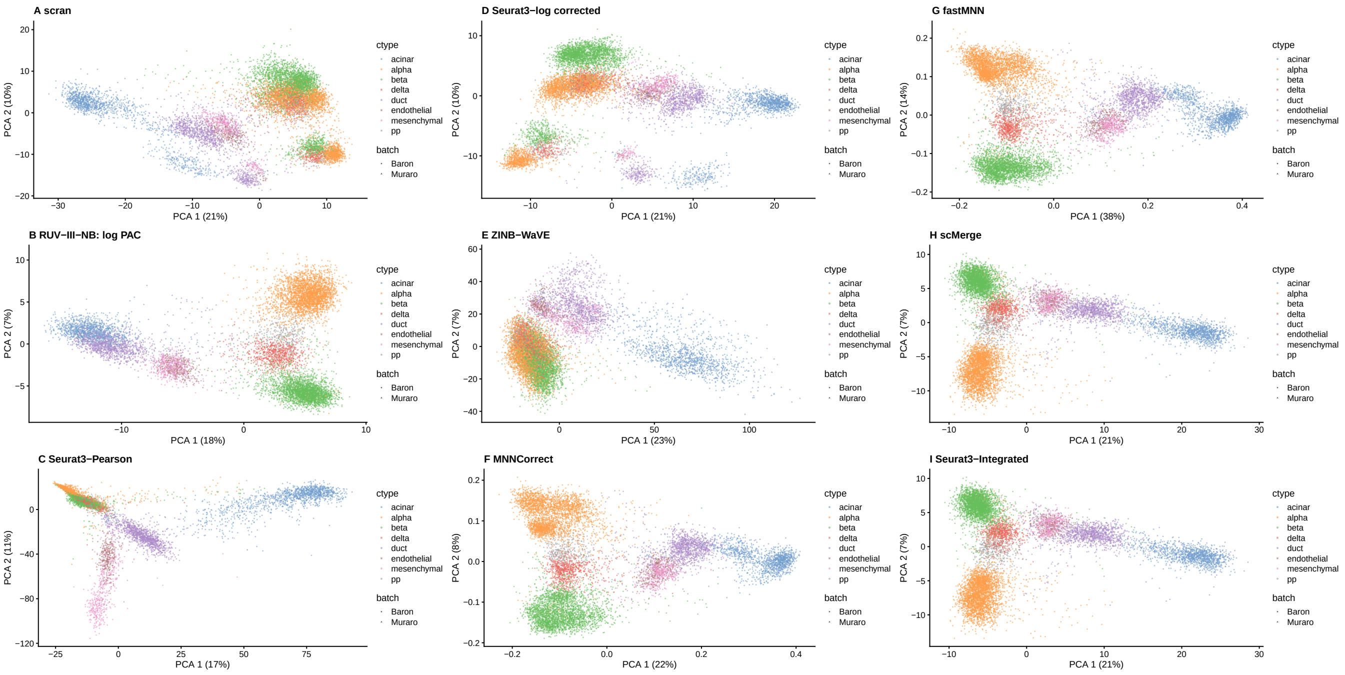


Supp. Fig. 11: Pancreas study. Leading PC of differently normalized counts. Coloured by cell type, while shape refers to batch.

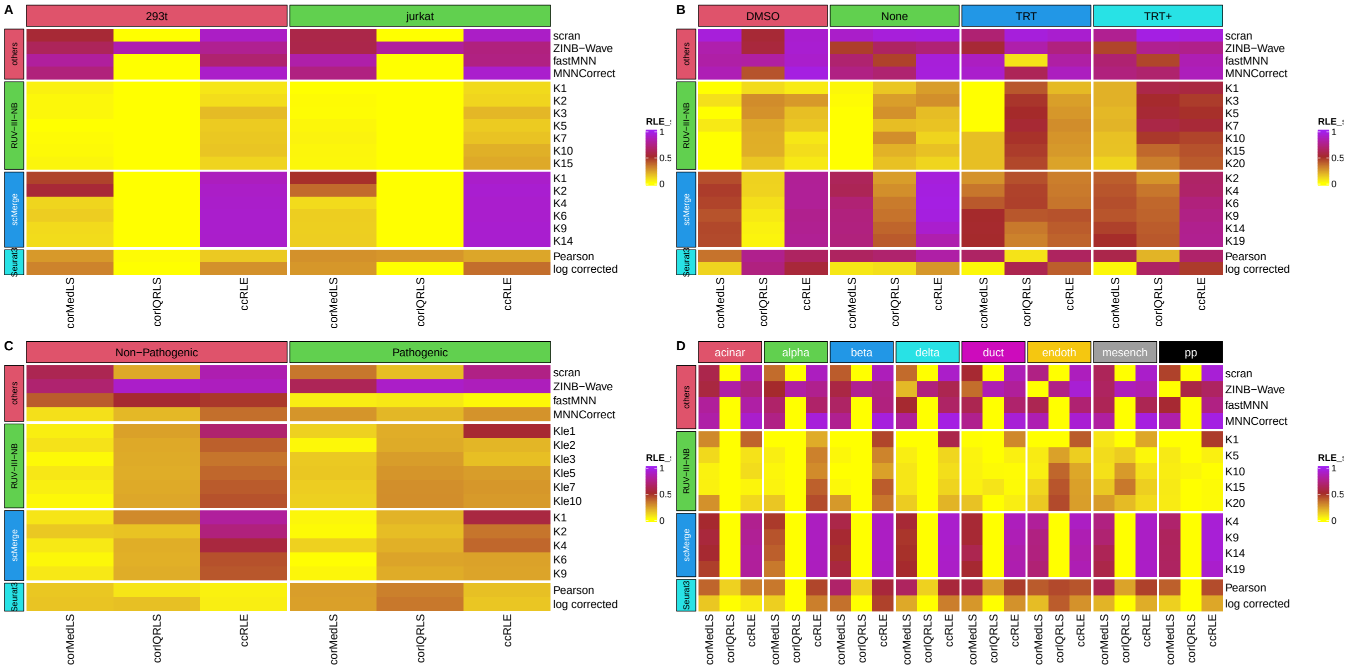


Supp. Fig. 12: Heatmap of correlations between RLE plot medians and log library sizes (first column), correlations between RLE plot IQR and log library sizes (second column) and canonical correlation between (RLE plot median, RLE plot IQR) and (log LS , batch variables) (third column), all stratified by cell type. (A) Cell line study. (B) CLL study. (C) Gaublomme study. (D) Pancreas study.

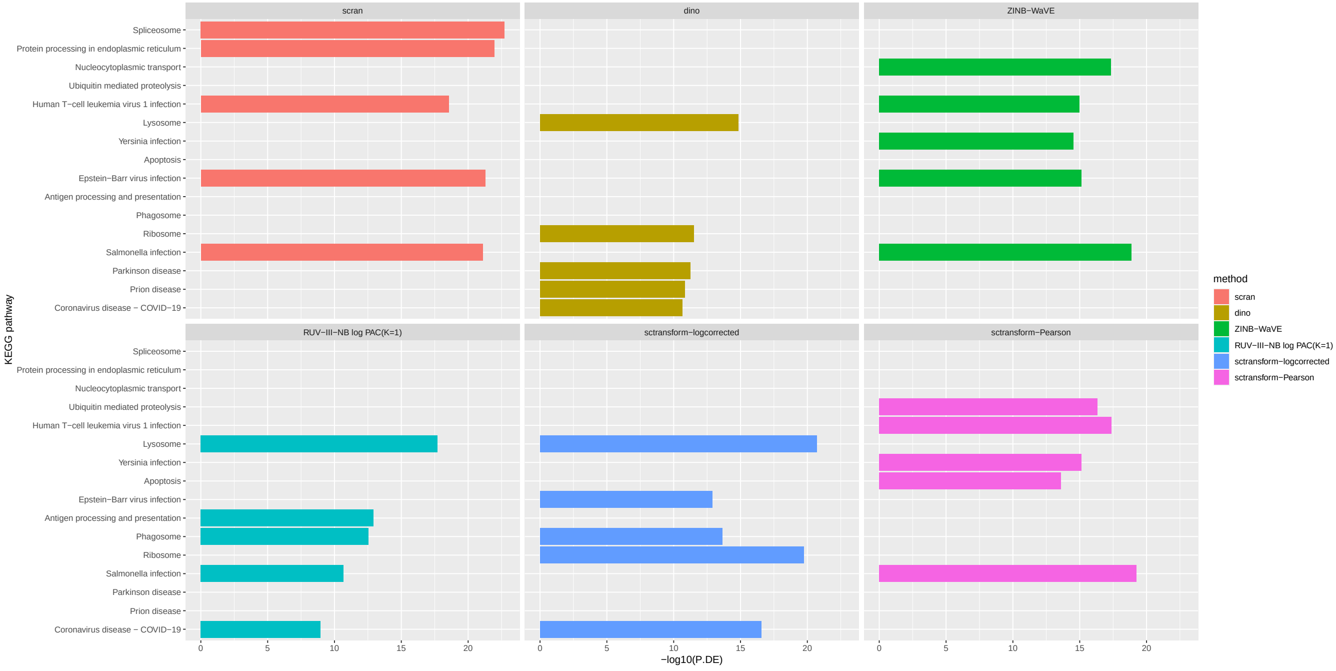


Supp. Fig. 13: Top 5 KEGG Pathways among DEG calculated from normalized data from the NSCLC study: larger vs smaller monocytes.

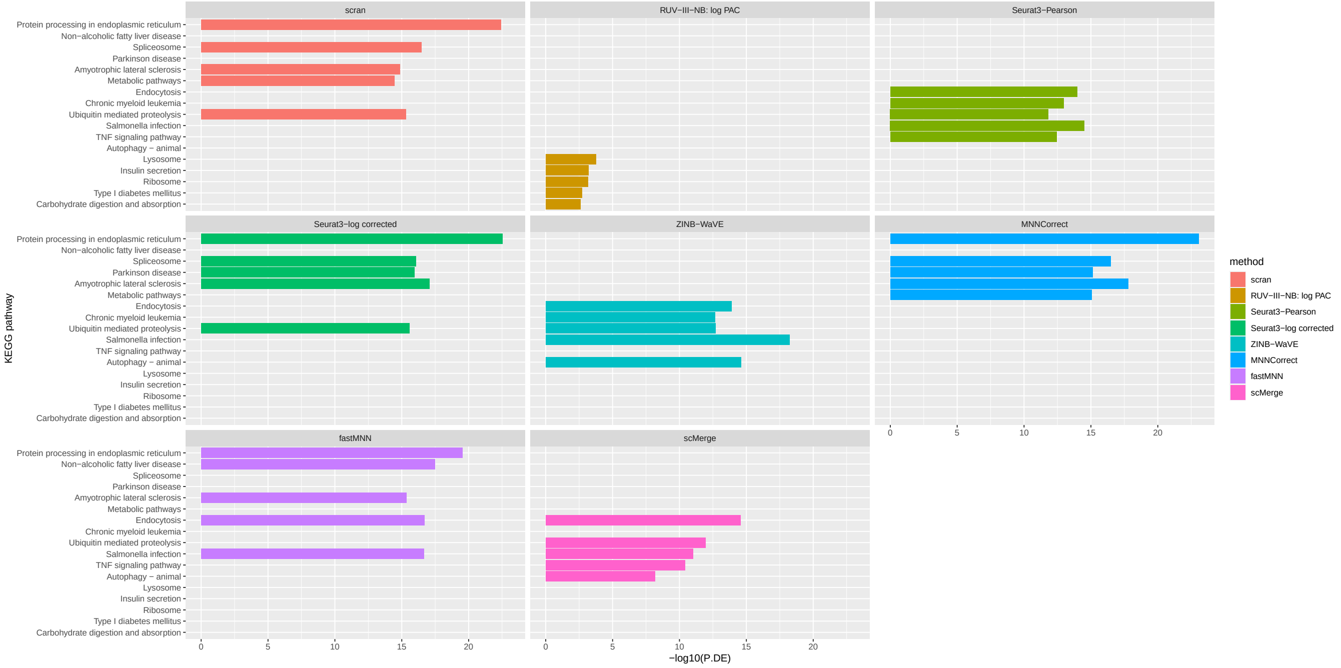


Supp. Fig. 14: Top 5 KEGG Pathways among DEG calculated from normalized data from the Pancreas study: larger vs smaller beta cells

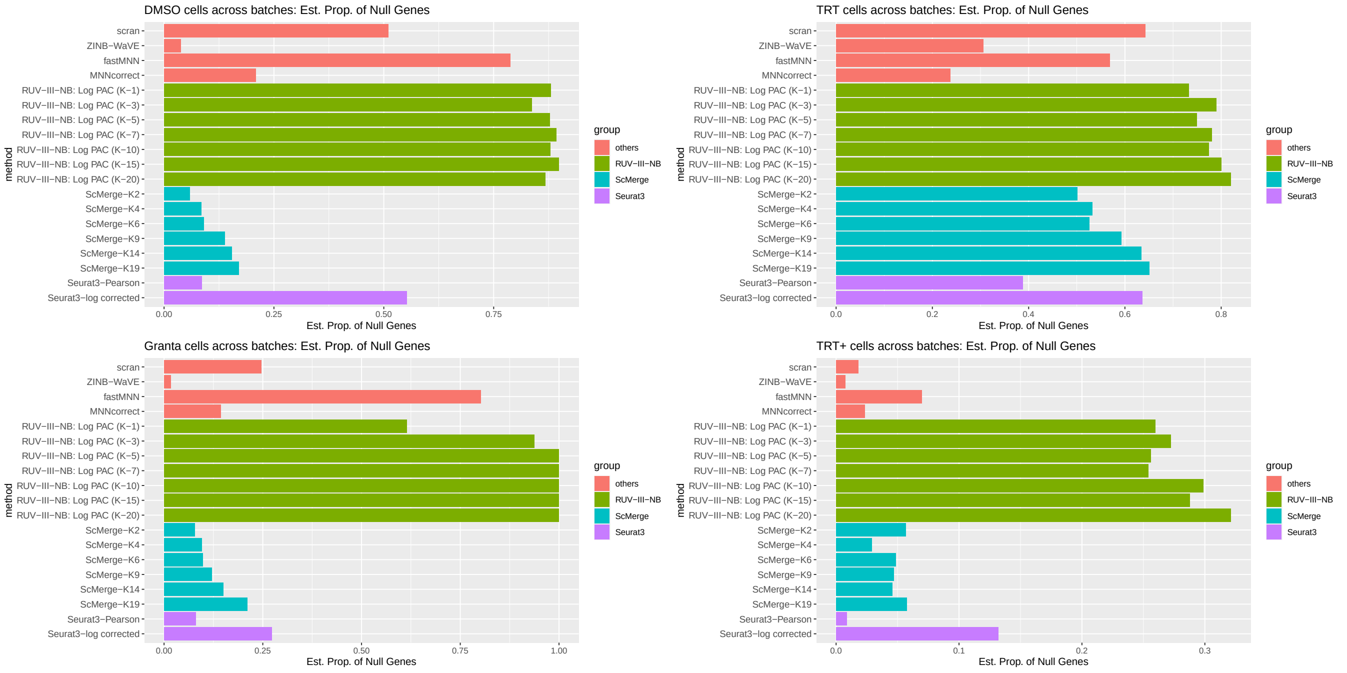


Supp. Fig. 15: CLL study. Estimated proportion of non-DE (i.e. null) genes when comparing the same cell type across batches. When batch effects are completely removed, the proportion of null genes is 1.

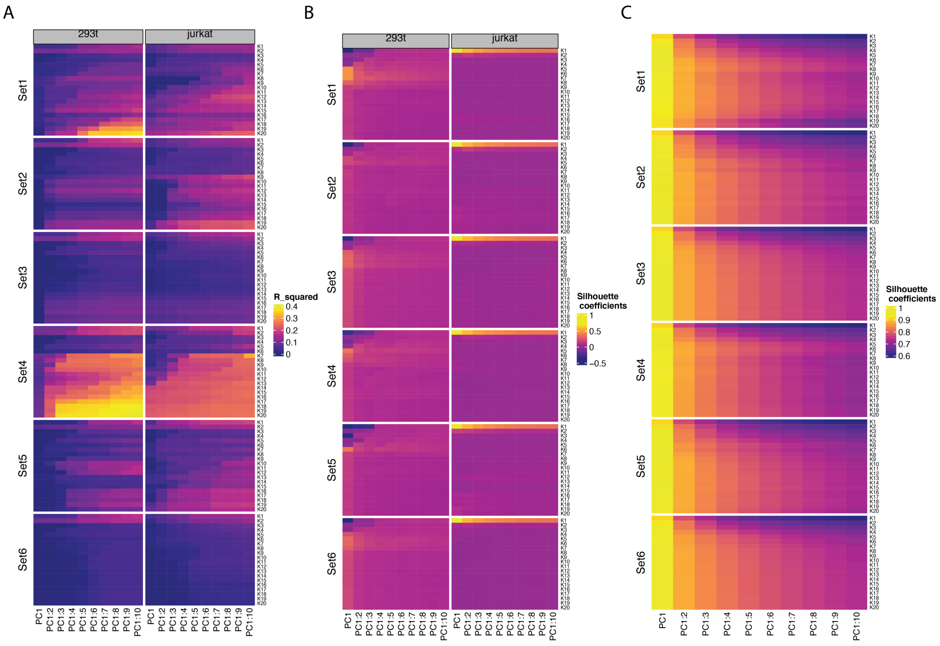


Supp. Fig. 16: Robustness of RUV-III-NB performance metrics across different negative control sets and numbers of unwanted factors ($K$) using the Cell line dataset. (A) $R^2$ between log library and cumulative principal components (PC). (B) Average Technical Silhouette width. (C) Average Biological Silhouette width. Set1: All genes are negative controls. Set2: scRNA-seq housekeeping genes from the scMerge package. Set3 : Use the scran::modelGeneVar function to select the 1000 genes with the highest technical variance in the batch with both cell lines (batch 3). Set4: DE analysis between cell lines in batch 3, followed by selecting the 1000 genes with the highest p-values. Set5: Use the scMerge::scSEGIndex function to select the 1000 genes with the most stable expression from batch 3. Set6: Use genes with biological absolute log fold-change $\leq 0.05$ (biological log fold-change is from DE analysis between the cell lines in batch 3) and technical absolute log fold-change $\geq 2$(technical log fold-change is from DE analysis between cell lines in batch 1 and 2).

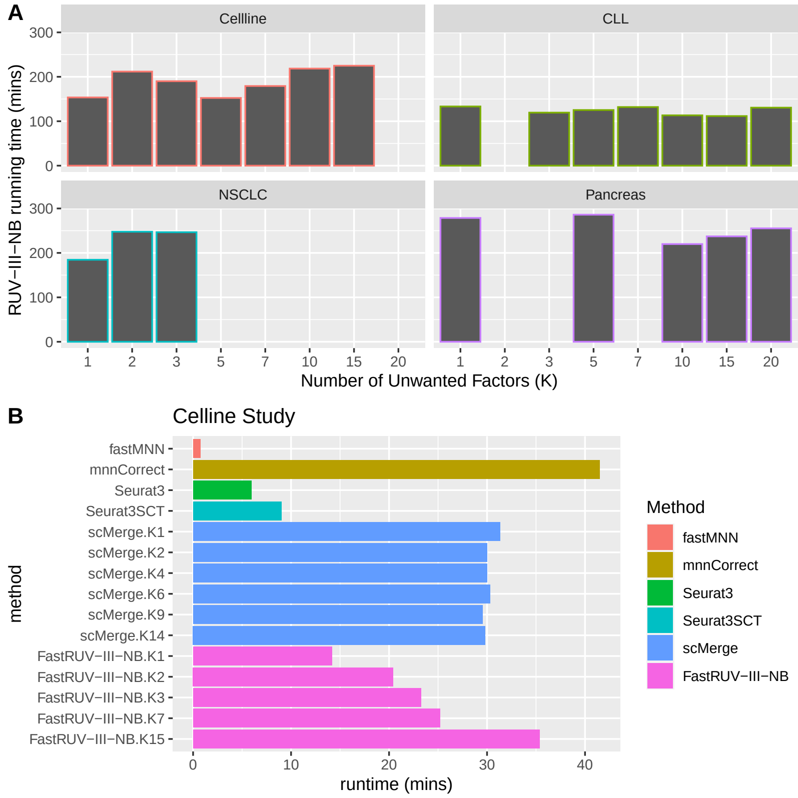


Supp. Fig. 17: (A) RUV-III-NB running time for various datasets on HPC environment with 15 cores and 120 Gb total RAM (8Gb RAM per core). (B) Comparison of Fast RUV-III-NB running time against other methods on a Linux PC with 6 cores and 24 Gb total RAM (4Gb RAM per core).

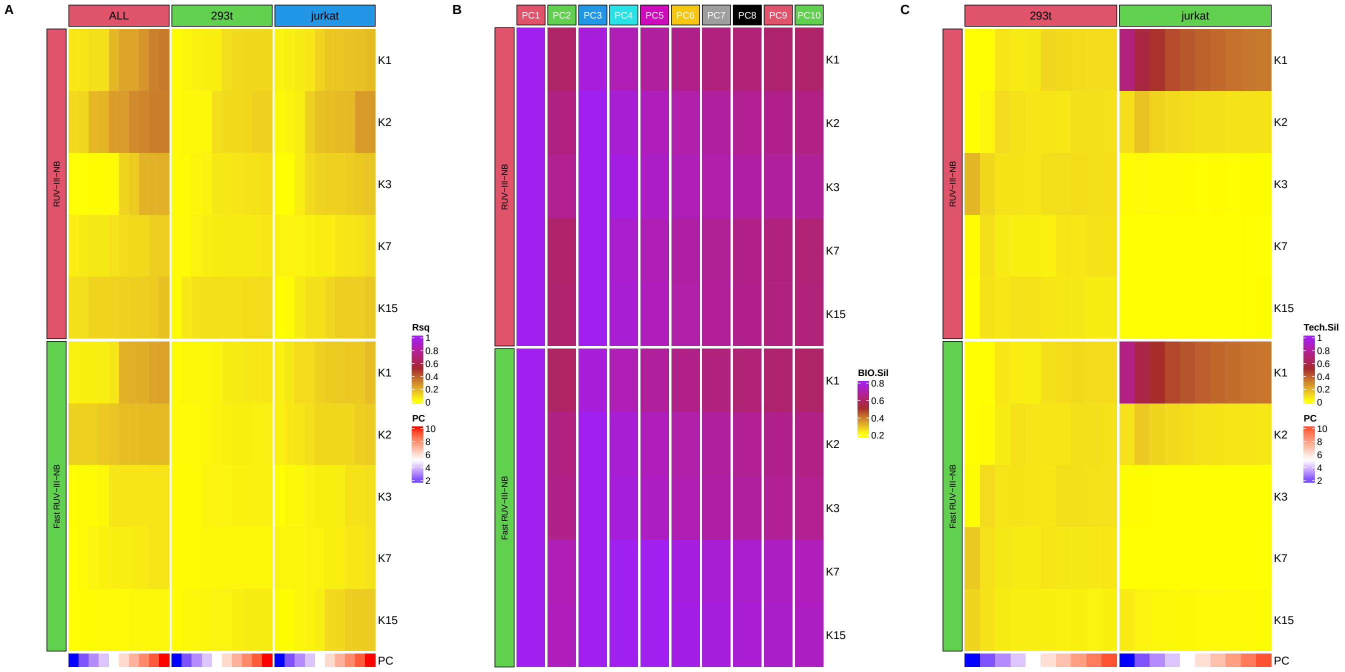


Supp. Fig. 18: Comparison of Fast vs standard RUV-III-NB implementation using the cell line study counts. (A) R-sq between leading PC and log library size. (B) Biological silhouette. (C). Technical (batch) silhouette.

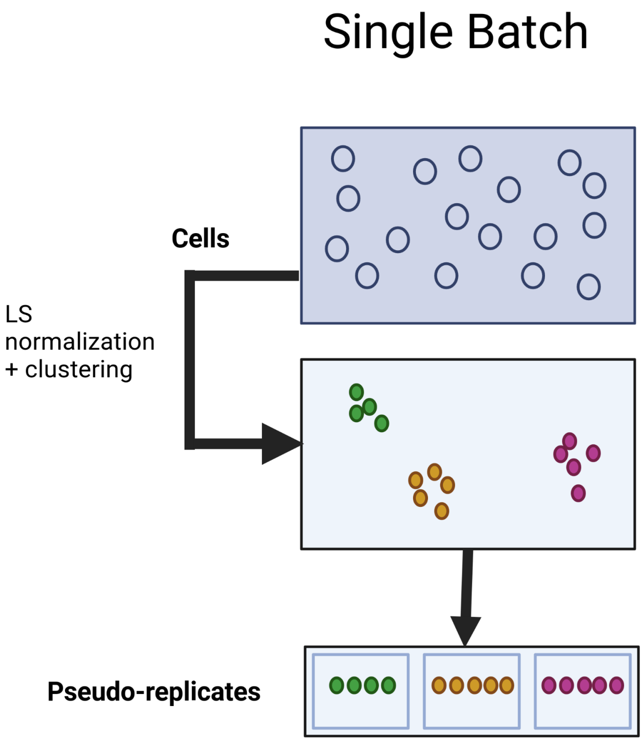


Supp. Fig. 19: Schematic illustration of RUV-III-NB pseudo-replicates identification with one batch. Cells are represented as circles and colour refers to biology.

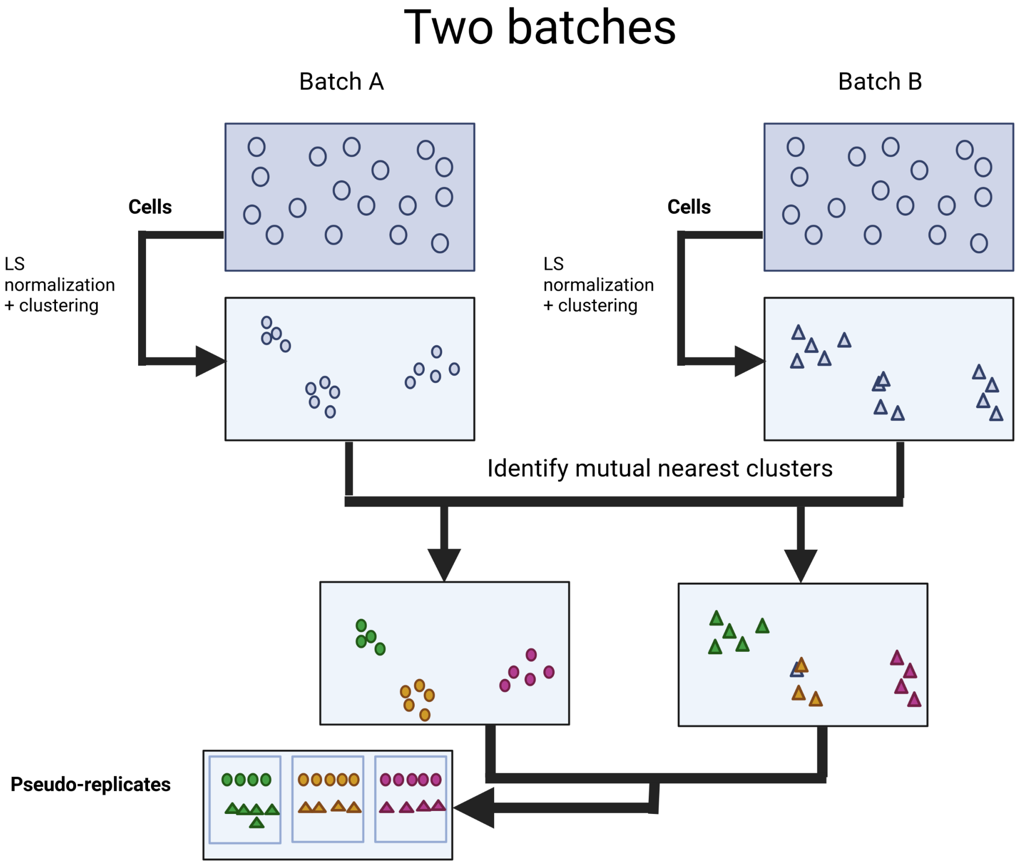


Supp. Fig. 20: Schematic illustration of RUV-III-NB pseudo-replicates identification with two batches. Cells from the first batch are represented as circles and those from the second batch are represented as triangles. Colour refers to biology.

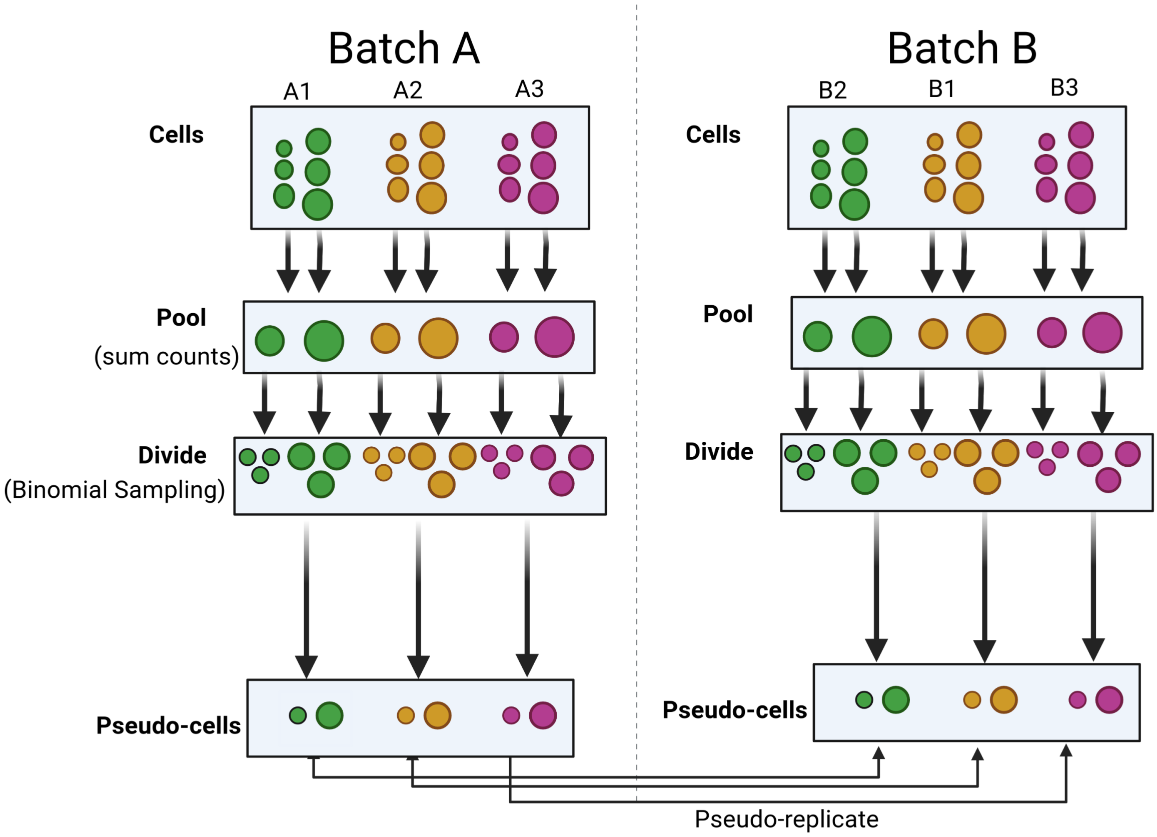


Supp. Fig. 21: Schematic illustration of RUV-III-NB pseudo-cells generation using pool-and-divide strategy with two batches. Cells are represented as circles with the size of the circles representing library size. Colour refers to biology. First, in each batch, cells of the same biology are split into $J = 2$ groups based on their library size. Then the cells from each group are pooled together, followed by Binomial sampling to divide the pooled counts equally and randomly selecting one of them as pseudo-cells. Pseudo-cells with the same biology are then declared as pseudo-replicates.
