## Supplementary Methods for "RUV-III-NB: Normalization of single cell RNA-seq Data"

Salim et al.

January 25, 2022

Let $\boldsymbol{y}_g = (y_{g1}, y_{g2}, \ldots, y_{gN})^T$ the vector of counts for gene $g$ across $N$ samples labelled by $c$. The RUV-III-NB model assumes that $y_{gc} \sim \text{NB}(\mu_{gc}, \psi_g)$, independently for different genes $g$ and different samples $c$. We will assume that there are *replicates* of the same sample among the $N$ samples, so that the number of *distinct samples* is $m < N$. In the context of scRNA-seq, the samples are single cells and replicate samples are cells belonging to the same sub-population that we call pseudo-replicates. The sub-populations can represent cell-types or another biological condition, while the equivalent of distinct samples in the single cell context will be called replicate sets, see main text for a fuller discussion.

For each gene $g$, we model $\boldsymbol{\mu}_g = (\mu_{g1}, \mu_{g2}, \ldots, \mu_{gN})^T$ as a function of unobserved unwanted factors $\boldsymbol{W}(N \times k)$ via the following generalized linear model (McCullagh and Nelder, 1989),

$$h(\boldsymbol{\mu}_g) = \zeta_g \mathbf{1} + \boldsymbol{M}\boldsymbol{\beta}_g + \boldsymbol{W}\boldsymbol{\alpha}_g \tag{1}$$

where $h(\boldsymbol{\mu}_g) = \log \boldsymbol{\mu}_g$ is the link function, $\boldsymbol{M}(N \times m)$ is the replicate membership matrix that links the $N$ cells to the $m$ distinct sub-populations, with $\boldsymbol{M}(i,j) = 1$ if the $i^{th}$ cell comes from the $j^{th}$ sub-population and 0 otherwise, $\boldsymbol{\beta}_g(m \times 1)$ is the vector of parameters that represent biological differences among the $m$ sub-populations, $\boldsymbol{\alpha}_g(k \times 1)$ is the vector of regression coefficient associated with the unwanted factors and finally $\zeta_g$ is location parameter for gene $g$ after adjusting for unwanted factors.

For a given $k$, our objective is to estimate $\boldsymbol{W}\boldsymbol{\alpha}_g$ and remove its effect from the counts. To do that we also need to estimate the other parameters, namely $\zeta_g$, $\boldsymbol{\beta}_g(m \times 1)$ and the dispersion parameter $\psi_g$.

#### Identifiability Issues

Note that in our model since both $\boldsymbol{W}$ and $\boldsymbol{\alpha}_g$ are unknown, without further constraints only the product $\boldsymbol{W}\boldsymbol{\alpha}_g$ is identifiable and not its component terms. To overcome this, we constrain each column of $\boldsymbol{W}$ to have unit norm, without requiring these columns to be orthogonal.

#### Negative control Genes

We also assume there is a set of negative control genes whose count variation across cells is representative of the unwanted variation that is to be removed, but is unrelated to the biology of interest. For these genes, $\boldsymbol{\beta}_g \approx 0$ and the model simplifies

$$h(\boldsymbol{\mu}_g) \approx \zeta_g \mathbf{1} + \boldsymbol{W}\boldsymbol{\alpha}_g \tag{2}$$

#### Shrinkage of $\boldsymbol{\alpha}_g$

Since our count data are often very sparse, we shrink the estimates of $\boldsymbol{\alpha}_g$ to stabilize them. This is achieved by treating $\boldsymbol{\alpha}_g$ as multivariate normal random variable, $\boldsymbol{\alpha}_g \sim MVN(\boldsymbol{\alpha}_\mu, \lambda_\alpha^{-1}\boldsymbol{I}_k)$ for a known $\lambda_\alpha$. Our model for gene $g$ can now be written

$$\begin{aligned} h(\boldsymbol{\mu}_g) &= \zeta_g \mathbf{1} + \boldsymbol{M}\boldsymbol{\beta}_g + \boldsymbol{W}\boldsymbol{\alpha}_g \\ &= \zeta_g \mathbf{1} + \boldsymbol{M}\boldsymbol{\beta}_g + \boldsymbol{W}\boldsymbol{\alpha}_\mu + \boldsymbol{W}\boldsymbol{\delta}_g \end{aligned}$$

where $\boldsymbol{\alpha}_g = \boldsymbol{\alpha}_\mu + \boldsymbol{\delta}_g$ , $\boldsymbol{\delta}_g \sim MVN(0, \lambda_\alpha^{-1} I_k)$ and $\lambda_\alpha$ is assumed to be known.

### RUV-III-NB Estimation via Modified Iteratively Reweighted Least Squares

#### Step 0: Initial parameter estimates

- We first obtain an initial estimate of $\boldsymbol{W}$ as the first $k$ right singular vector of $\log(\boldsymbol{Y_c^*}+1)$, where $\boldsymbol{Y_c^*}(G_c \times N)$ is the matrix of sequencing counts for $G_c$ negative control genes that has been scaled gene-by-gene by gene-specific average count.

- Given the initial estimate of $\boldsymbol{W}$, we set the initial estimates of $\beta_{\mathbf{g}}$=0 and obtain the initial estimates of $\boldsymbol{\zeta_g}$ and $\boldsymbol{\alpha}_g$ using weighted linear regression of $\log(\boldsymbol{y_g}+1)$ on the initial estimate of $\boldsymbol{W}$. We use $\boldsymbol{y_g}+1$ as weight so that observations with higher mean count are assigned larger weights, just like in Poisson regression.

- The initial estimate of $\boldsymbol{\alpha_\mu}$ is calculated as the average of the initial estimate of $\boldsymbol{\alpha_g}$ across genes, and the initial estimate of $\boldsymbol{\delta_g}$ is calculated simply as the deviation of the initial estimate of $\boldsymbol{\alpha_g}$ from the mean.

- Obtain initial estimates of the dispersion parameter, $\psi_g^0$ using `edgeR::estimateDisp` (Robinson *et al.*, 2010) function with the initial estimate of $\boldsymbol{W\alpha_g}$ as offset term.

- We call all these estimates, "current" estimates and use the modified iteratively reweighted least squares (IRLS) algorithm below to improve the estimates until convergence is achieved

For the initial dispersion parameter estimate $\psi_g^0$, in each IRLS iteration, given the current parameter estimates $\boldsymbol{\alpha}_\mu^r(k\times 1)$, $\boldsymbol{\delta}_g^r(k\times 1)$ and $\boldsymbol{\beta}_g^r(m\times 1)$, we use the first-order Taylor approximation to (1) to obtain the working linear model

$$\boldsymbol{z}_g^r = \zeta_g^r\mathbf{1} + \boldsymbol{M}\boldsymbol{\beta}_g^r + \boldsymbol{W}^r\boldsymbol{\alpha}_\mu^r + \boldsymbol{W}^r\boldsymbol{\delta}_g^r + \boldsymbol{e}_g^r \tag{3}$$

where $\boldsymbol{z}_g^r$ is the current working vector, $\boldsymbol{\mu}_g^r = \exp(\zeta_g^r\mathbf{1} + \boldsymbol{M}\boldsymbol{\beta}_g^r + \boldsymbol{W}^r\boldsymbol{\alpha}_\mu^r + \boldsymbol{W}^r\boldsymbol{\delta}_g^r)$ and $\boldsymbol{e}_g^r = \frac{\boldsymbol{y}_g-\boldsymbol{\mu}_g^r}{\boldsymbol{\mu}_g^r}$ is the vector of current working residuals, with variance for its $c$th component approximately equal to,

$$\text{var}(e_{gc}^r) \approx \text{var}\Big(\frac{y_{gc}}{\mu_{gc}^r}\Big) = \frac{1}{\mu_{gc}^r} + \psi_g^0. \tag{4}$$

Hence, across all observations

$$\text{var}(\boldsymbol{e}_g^r) = \text{Diag}(1/\mu_{gc}^r + \psi_g^0) = \mathbf{\Sigma_g^r}.$$

Equation (3) is for a specific gene $g$. When writing models for all genes $g = 1,\dots G$, it will be convenient to put them in matrix form as

$$\boldsymbol{Z}^r = \mathbf{1}\boldsymbol{\zeta}^{rT} + \boldsymbol{M}\boldsymbol{\beta}^r + \boldsymbol{W}^r\boldsymbol{\alpha}_\mu^r + \boldsymbol{W}^r\boldsymbol{\delta}^r + \boldsymbol{e}^r. \tag{5}$$

Here, $\boldsymbol{Z}^r(N\times G)$ has a column vector for each gene, and similarly, $\boldsymbol{\beta}^r(m\times G)$, $\boldsymbol{\delta}^r(k\times G)$ and $\boldsymbol{e}^r(N\times G)$ are matrices with $\boldsymbol{\beta}_g^r$, $\boldsymbol{\delta}_g^r$ and $\boldsymbol{e}_g^r$ as columns, while $\boldsymbol{\zeta}^r(G\times 1)$ is a vector with $\zeta_g^r$ as its elements.

For a given $\mathbf{W^r}$, equation (3) is a (normal) linear regression model with unequal (heteroscedastic) variances. The log-likelihood after incorporating the shrinkage penalty for $\boldsymbol{\alpha}_g$ can be written as follows

$$-\sum_{g=1}^{G}(\boldsymbol{z}_g^r - \zeta_g^r\mathbf{1} - \boldsymbol{M}\boldsymbol{\beta}_g^r - \boldsymbol{W}^r\boldsymbol{\alpha}_\mu^r - \boldsymbol{W}^r\boldsymbol{\delta}_g^r)'\mathbf{\Omega}_g^r(\boldsymbol{z}_g^r - \zeta_g^r\mathbf{1} - \boldsymbol{M}\boldsymbol{\beta}_g^r - \boldsymbol{W}^r\boldsymbol{\alpha}_\mu^r - \boldsymbol{W}^r\boldsymbol{\delta}_g^r) + \lambda_\alpha\sum_{g=1}^{G}\boldsymbol{\delta}_g'\boldsymbol{\delta}_g \tag{6}$$

where $\mathbf{\Omega_g^r}$ is the inverse of $\mathbf{\Sigma_g^r}$ with main diagonal elements $(1/\mu_{gc}^r + \psi_g^0)^{-1}$.

If $\boldsymbol{W}$ had been known, estimating the other parameters would have been easily done by standard IRLS. But since $\boldsymbol{W}$ also needs to be estimated, we need to use the following variant IRLS that takes advantage of the control genes and the replicate matrix $\boldsymbol{M}$ to estimate $\boldsymbol{W}$ and its regression coefficients $\boldsymbol{\alpha}_g$. In its essence, it is an adaptation to the present GLM context of the linear model version of RUV-III.

- **Step 1:** Calculate the current working vector $z_g^r$ and apply the residual operator $R_M = \boldsymbol{I} - \boldsymbol{M}(\boldsymbol{M^T M})^{-\mathbf{1}}\boldsymbol{M^T}$ to the working vector so that model in equation (3) can be written as

$$R_M\boldsymbol{z}_g^r \;\; = \;\; R_M\boldsymbol{W}^r\boldsymbol{\alpha}_\mu^r + R_M\boldsymbol{W}^r\boldsymbol{\delta}_g^r + R_M\boldsymbol{e}_g.$$

The log-likelihood for this new data is

$$-\sum_{g=1}^{G} R_M(\boldsymbol{z_g^r} - \boldsymbol{W^r}\boldsymbol{\alpha_\mu^r} - \boldsymbol{W^r}\boldsymbol{\delta_g^r})'\boldsymbol{\Omega}_g^r R_M(\boldsymbol{z_g^r} - \boldsymbol{W^r}\boldsymbol{\alpha_\mu^r} - \boldsymbol{W^r}\boldsymbol{\delta_g^r}) + \lambda_\alpha \sum_{g=1}^{G} \boldsymbol{\delta}_g'\boldsymbol{\delta}_g. \quad (7)$$

- **Step 2:** Using the model for the transformed working vector, we update the estimates of $\boldsymbol{\alpha}_\mu$ and $\boldsymbol{\delta}_g$ by differentiating the log-likelihood above w.r.t to $\boldsymbol{\alpha}_\mu^r$ and $\boldsymbol{\delta}_g^r$ and setting them to zero. This gives the following update formulae

$$\boldsymbol{\alpha}_\mu^{r+1} = \left(\sum_{g=1}^{G} \boldsymbol{W^{r\prime}} R_M \boldsymbol{\Omega_g^r} R_M \boldsymbol{W^r}\right)^{-1} \left(\sum_{g=1}^{G} \boldsymbol{W^{r\prime}} R_M \boldsymbol{\Omega_g^r} R_M (\boldsymbol{z_g^r} - R_M \boldsymbol{W^r}\boldsymbol{\delta}_g^r)\right) \quad (8)$$

$$\boldsymbol{\delta}_g^{r+1} = (\boldsymbol{W^{r\prime}} R_M \boldsymbol{\Omega_g^r} R_M \boldsymbol{W^r} + \lambda_\alpha \boldsymbol{I}_k)^{-1} \boldsymbol{W^{r\prime}} R_M \boldsymbol{\Omega_g^r} R_M (\boldsymbol{z_g^r} - R_M \boldsymbol{W^r}\boldsymbol{\alpha}_\mu^{r+1}) \quad (9)$$

- **Step 3:** To update $\boldsymbol{W^r} \to \boldsymbol{W^{r+1}}$, we take advantage of the negative control genes. Note that by definition of these genes, we have $\boldsymbol{\beta}_g \approx 0$. If we denote the current working vector for cell $c$ associated with negative control gene subset $C$ as $\boldsymbol{z}_{C,c}^r(G_c \times 1)$, we have

$$\boldsymbol{z}_{C,c}^r - \boldsymbol{\zeta}_C^r = \boldsymbol{\alpha}_c^{r\prime} w_c^r + \boldsymbol{e}_{C,c}^r$$

where $\boldsymbol{\zeta}_C^r(G_c \times 1)$ is the subset of vector $\boldsymbol{\zeta}^r$ associated with control genes and $\boldsymbol{\alpha}_C^r(k \times G_c)$ is the subset of matrix $\boldsymbol{\alpha^r} = \boldsymbol{\delta^r} + \boldsymbol{\alpha_\mu^r}\boldsymbol{1^T}$, whose columns associated with the control genes, $\boldsymbol{w_c^r}(k \times 1)$ the row of $\boldsymbol{W}^r$ associated with cell $c$.

An updated estimate $\boldsymbol{w_c}$ can then be obtained as

$$\boldsymbol{w_c^{r+1}} = (\boldsymbol{\alpha}_C^r \boldsymbol{\Omega}_{C,c}^r \boldsymbol{\alpha}_C^{r\prime})^{-1} \boldsymbol{\alpha}_C^r \Omega_{C,c}^r (\boldsymbol{z}_{C,c}^r - \boldsymbol{\zeta}_C^r)$$

where $\boldsymbol{\Omega}_{C,c}^r(G_c \times G_c)$ is a diagonal matrix with main diagonal elements equal to the inverse of current variance estimate of $\boldsymbol{z}_{C,c}^r$.

- **Step 4:** To update $\zeta_g^r \to \zeta_g^{r+1}$, we differentiate equation (6) w.r.t $\zeta_g^r$ and set it to zero, to obtain the update formula

$$\zeta_g^{r+1} = (\boldsymbol{1'}\boldsymbol{\Omega_g^r}\boldsymbol{1})^{-1}\boldsymbol{1'}\boldsymbol{\Omega_g^r}(\boldsymbol{z_g^r} - \boldsymbol{M}\beta^r - \boldsymbol{W^r}\boldsymbol{\alpha}_\mu^r - \boldsymbol{W^r}\boldsymbol{\delta}_g^r)$$

- **Step 5:** The last step involves updating $\boldsymbol{\beta}_g^r$, done in the following way. First, note the part of equation (6) that involves $\boldsymbol{\beta}_g^r$ is given by

$$(\boldsymbol{z_g^r} - \zeta_g^r\boldsymbol{1} - \boldsymbol{M}\boldsymbol{\beta}_g^r - \boldsymbol{W^r}\boldsymbol{\alpha}_\mu^r - \boldsymbol{W^r}\boldsymbol{\delta}_g^r)'\boldsymbol{\Omega_g^r}(\boldsymbol{z_g^r} - \zeta_g^r\boldsymbol{1} - \boldsymbol{M}\boldsymbol{\beta}_g^r - \boldsymbol{W^r}\boldsymbol{\alpha}_\mu^r - \boldsymbol{W^r}\boldsymbol{\delta}_g^r)$$

If we differentiate the log-likelihood w.r.t $\boldsymbol{\beta}_g^r$ and set it to zero, we obtain the update formula

$$\boldsymbol{\beta}_g^{r+1} = (\boldsymbol{M'}\boldsymbol{\Omega_g^r}\boldsymbol{M})^{-1}\boldsymbol{M'}\boldsymbol{\Omega_g^r}(\boldsymbol{z_g^r} - \zeta_g^r\boldsymbol{1} - \boldsymbol{W^r}\boldsymbol{\alpha}_\mu^r - \boldsymbol{W^r}\boldsymbol{\delta}_g^r)$$

**Note:** the above formula involves matrix inversion for each of potentially thousands of genes. But in practice we did not perform the time-consuming matrix inversion. This is because due to the structure of matrix $\boldsymbol{M}$, it can be shown that for each unique sample, the above update formula will reduce to weighted average of the corrected working vector $\boldsymbol{z_g^{*r}} = (\boldsymbol{z_g^r} - \zeta_g^r\boldsymbol{1} - \boldsymbol{W^r}\boldsymbol{\alpha}_\mu^r - \boldsymbol{W^r}\boldsymbol{\delta}_g^r)$ with the main diagonal elements of $\boldsymbol{\Omega}_g^r$ as weights. For each distinct sample, the weighted average is taken across elements of the corrected working vector that correspond to the sub-population of that cell. For singleton cells this will simply equal to the element of the corrected working vector associated with that cell.

**Shrinkage of $\boldsymbol{\beta}_g$:** To estimate $\boldsymbol{\beta}_g$ we have to estimate $m$ parameters for each gene, giving a total of $G \times m$ parameters across all genes. We found that it is often necessary to regularize this process by treating $\boldsymbol{\beta}_g$ as a multivariate normal random variable, $\boldsymbol{\beta}_g \sim N_m(0, \lambda_\beta^{-1}\boldsymbol{I}_m)$ with known $\lambda_\beta$. Under this assumption, the update formula is given by

$$\beta_g^{r+1} = (\boldsymbol{M}'\boldsymbol{\Omega}_{\boldsymbol{g}}^r\boldsymbol{M} + \lambda_\beta \boldsymbol{I}_m)^{-1}\boldsymbol{M}'\boldsymbol{\Omega}_{\boldsymbol{g}}^r(\boldsymbol{z}_{\boldsymbol{g}}^r - \zeta_g^r\mathbf{1} - \boldsymbol{W}^r\boldsymbol{\alpha}_\mu^r - \boldsymbol{W}^r\boldsymbol{\delta}_g^r)$$

This regularized estimate will be shrunken towards zero compared to the non-regularized estimate. The amount of shrinkage will vary across genes. More abundant genes have larger mean parameters and main diagonal elements of $\boldsymbol{\Omega}_g^r$ and so will be shrunk less, while estimates for the less abundant genes will be shrunk more.

- **Step 6:** Iterate between Steps 1 and 5 until convergence (determined using deviance) is achieved for $\boldsymbol{W}, \boldsymbol{\alpha}, \boldsymbol{\beta}$. Then, update the dispersion parameter, gene-by-gene, $\psi_g^0 \rightarrow \psi_g^1$. This step is achieved using `edgeR::estimateDisp` function from `edgeR` package that employs regularization on the dispersion parameter estimates (Robinson *et al.*, 2010).

- **Step 7:** Stop when overall convergence is achieved; otherwise go to step 1 and re-estimate the parameters using the updated estimates of the dispersion parameters.

### Robust Weights

By default during IRLS, we use $\omega_{gc}^r = (1/\mu_{gc}^r + \psi_g^0)^{-1}$ as an observation weight. This weight can be robustified using the Huber weight function. In the robust version, we use $\omega_{gc}^{*r} = \omega_{gc}^r \times \psi(d_{gc}^r)$, where $\psi(.)$ is the Huber weight function and $d_{gc}^r$ is the signed deviance given by

$$\text{sign}(y_{gc} - \hat{\mu}_{gc}^r)\sqrt{2\{\log \ell(\mu_{gc} = y_{gc}, \psi_g; y_{gc}) - \log \ell(\mu_{gc} = \hat{\mu}_{gc}^r, \psi_g; y_{gc})\}}$$

where $\hat{\mu}_{gc}^r$ is the current estimates of NB mean parameter for gene $g$ and cell $j$.

### Adjusted Data

Upon convergence, we obtain estimates of the unwanted variation $\hat{\boldsymbol{W}}\hat{\boldsymbol{\alpha}}_{\boldsymbol{g}}$. Then, their effects need to removed from the counts after which downstream analyses such as clustering, trajectory and differential expression analyses can be performed. RUV-III-NB provides two forms of adjusted data:

- Pearson residuals:

$$\frac{y_{gc} - \hat{\mu}_{gc}}{\sqrt{\hat{\mu}_{gc} + \hat{\mu}_{gc}^2\hat{\psi}_g^2}}$$

  where $\hat{\mu}_{gc} = \exp(\hat{\zeta}_g + \hat{\boldsymbol{w}}_{\boldsymbol{c}}{}^{\boldsymbol{T}}\hat{\boldsymbol{\alpha}}_g)$ and $\hat{\boldsymbol{w}}_{\boldsymbol{c}}$ the c$^{th}$ row of the matrix $\hat{\boldsymbol{W}}$.

  When $k = 1$ and $\hat{\mathbf{W}}$ is approximately equal to log library size (up to a scaling factor), these Pearson residuals will roughly agree with those of (Hafemeister and Satija, 2019), with different shrinkage of parameter estimates leading to small differences. When $k > 1$ and some columns of $\mathbf{W}$ reflect batch effects, these Pearson residuals will also adjust for unwanted variation other than library size, such as batch effects.

- Log of percentile-invariant adjusted count (PAC):

$$\log(F^{-1}(r_{gc}; \mu_{gc} = \exp(\hat{\zeta}_g + \boldsymbol{m}_{\boldsymbol{c}}^{\boldsymbol{T}}\hat{\boldsymbol{\beta}}_{\boldsymbol{g}} + \bar{\boldsymbol{w}}^{\boldsymbol{T}}\hat{\boldsymbol{\alpha}}_{\boldsymbol{g}}), \hat{\psi}_g) + 1)$$

  where $r_{gc} \sim U(a_{gc}, b_{gc})$ and

$$\begin{aligned} a_{gc} &= F(y_{gc}; \mu_{gc} = \exp(\hat{\zeta}_g + \boldsymbol{m}_{\boldsymbol{c}}^{\boldsymbol{T}}\hat{\boldsymbol{\beta}}_{\boldsymbol{g}} + \hat{\boldsymbol{w}}_{\boldsymbol{c}}{}^{\boldsymbol{T}}\hat{\boldsymbol{\alpha}}_{\boldsymbol{g}}, \hat{\psi}_g)) \\ b_{gc} &= F(y_{gc} + 1; \mu_{gc} = \exp(\hat{\zeta}_g + \boldsymbol{m}_{\boldsymbol{c}}^{\boldsymbol{T}}\hat{\boldsymbol{\beta}}_{\boldsymbol{g}} + \hat{\boldsymbol{w}}_{\boldsymbol{c}}{}^{\boldsymbol{T}}\hat{\boldsymbol{\alpha}}_{\boldsymbol{g}}, \hat{\psi}_g)) \end{aligned}$$

  where $F(.)$ is the negative binomial c.d.f and $F^{-1}(.)$ its inverse, $\boldsymbol{m}_c$ is the c$^{th}$ row of the matrix $\boldsymbol{M}$, $\hat{\boldsymbol{w}}_{\boldsymbol{c}}$ the c$^{th}$ row of the matrix $\hat{\boldsymbol{W}}$ and $\bar{\boldsymbol{w}}$ is vector of entries equal to the average level $N^{-1}\sum_{c=1}^{N}\hat{\boldsymbol{w}}_{\boldsymbol{c}}$ of unwanted variation. Here $U(a, b)$ denoted a random variable uniformly distributed over the interval $(a, b)$.

### ZINB extension

The negative binomial (NB) model works well for scRNA-seq data with UMI. Like (Cao *et al.*, 2021), we found that scRNA-seq data without UMI contains excess zero counts relative to the NB distribution. Because of this, for non-UMI read counts, we extended the above model to Zero Inflated Negative Binomial (ZINB) model. The full specification of the ZINB model is as follows:

$$
\begin{aligned}
Y^*_{gc} &\sim ZINB(\pi^0_{gc}, \mu^*_{gc}, \psi^*_g) \\
\log \mu^*_{gc} &= \zeta_g + m_c^T \beta_g + w_{1c}\alpha^*_{g1} + \sum_{k=2}^{K} w_{kc}\alpha_{gk}
\end{aligned} \tag{10}
$$

where we use $Y^*_{gc}$ to denote random variable that represents the non-UMI read count and differentiates it from the random variables $Y_{gc}$ representing UMI counts.

Let us suppose we have two cells with the same library size and other unwanted factors, the same underlying cell-type (biology) *but have been sequenced using two different assays, one using UMI and the other giving non-UMI read counts.* Intuitively, these two cells must have the same expected counts for all genes,

$$
\begin{aligned}
E(Y_{gc}) &= E(Y^*_{gc}) \\
\mu_{gc} &= (1-\pi^0_{gc})\mu^*_{gc} \\
\log \mu_{gc} &= \log(1-\pi^0_g) + \log \mu^*_{gc}
\end{aligned}
$$

Substituting in equation 10 and using the fact that for UMI counts we can write the model for the mean as

$$
\log \mu_{gc} = \zeta_g + m_c^T \beta_g + w_{1c}\alpha_{g1} + \sum_{k=2}^{K} w_{kc}\alpha_{gk}
$$

we have

$$
\begin{aligned}
\zeta_g + m_c^T \beta_g + w_{1c}\alpha_{g1} + \sum_{k=2}^{K} w_{kc}\alpha_{gk} &= \log(1-\pi^0_{gc}) + \zeta_g + m_c^T \beta_g + w_{1c}\alpha^*_{g1} + \sum_{k=2}^{K} w_{kc}\alpha_{gk} \\
w_{1c}\alpha_{g1} &= w_{1c}\alpha^*_{g1} + \log(1-\pi^0_{gc}) \\
(\alpha^*_{g1} - \alpha_{g1}) &= -\log(1-\pi^0_{gc}) \\
\alpha^*_{g1} &= \alpha_{g1} - \log(1-\pi^0_{gc}).
\end{aligned}
$$

If instead of one cell we have $N$ cells, we would require $\sum_{c=1}^{N} E(Y_{gc}) = \sum_{c=1}^{N} E(Y^*_{gc})$ which leads to the following relationship between $\alpha_{g1}$ and $\alpha^*_{g1}$,

$$
\alpha^*_{g1} = \alpha_{g1} - \frac{\sum_{c=1}^{N} \log(1-\pi^0_{gc})}{\sum_c w_{1c}}
$$

We conclude that apart from the addition of zero-inflation parameter $\pi^0_{gc}$, only the regression coefficient associated with the first unwanted factor is different between the NB and ZINB models.

#### ZINB parameter estimation

Parameter estimation for ZINB model can be carried out using the same modified IWLS algorithm, with two important differences:

- Because the ZINB model has one extra parameter, we need an extra step (step 5a) where we update the zero-inflation parameter estimate $\pi^0_{gc}$ by maximizing the ZINB log-likelihood w.r.t to $\pi^0_{gc}$ with the other parameter values set at their current estimates.
- Because a ZINB is essentially a mixture of an NB distribution with a probability mass at zero, the estimation of the parameters for the NB components, i.e. the regression parameters for the mean and the dispersion parameter needs to use an *observation weight* that gets included in the IWLS (and robust)

weight. This observation weight is the conditional probability that the observation belongs to the NB component of the ZINB distribution and it is 1 for all non-zero counts. For zero counts this probability is given by

$$\frac{(1-\pi^0_{gc})f_{NB}(0;\mu=\mu^*_{gc},\psi=\psi^*_g)}{\pi^0_{gc}+(1-\pi^0_{gc})f_{NB}(0;\mu=\mu^*_{gc},\psi=\psi^*_g)}$$

When $\pi^0_{gc}=0$, the ZINB model reduces to the NB and the observation weights for all observation including the zero counts above will all equal to 1.

#### Adjusted Data under the ZINB model

- Pearson residuals:

$$\frac{y^*_{gc}-\hat{\mu}^*_{gc}(1-\hat{\pi}^0_{gc})}{\sqrt{\hat{\mu}^*_{gc}(1-\hat{\pi}^0_{gc})(1+\hat{\mu}^*_{gc}(\hat{\pi}^0_{gc}+\hat{\psi}^*_g))}}$$

  where $\hat{\mu}^*_{gc}=\exp(\hat{\zeta}_g+\hat{\boldsymbol{w}}_{\boldsymbol{c}}^{\boldsymbol{T}}\hat{\boldsymbol{\alpha}}^*_g)$.

- Log of percentile-invariant adjusted count:

$$\log(F^{-1}(r_{gc};\mu_{gc}=\exp(\hat{\zeta}_g+\boldsymbol{m}_c^T\hat{\boldsymbol{\beta}}_{\boldsymbol{g}}+\bar{\boldsymbol{w}}^T\hat{\boldsymbol{\alpha}}^*_{\boldsymbol{g}}),\psi=\hat{\psi}^*_g,\pi^0=\bar{\pi}^0_g)+1)$$

  where $r_{gc}\sim U(a_{gc},b_{gc})$ is the randomized quantile (Dunn and Smyth, 1996)

$$\begin{aligned}a_{gc} &= F(y_{gc};\mu_{gc}=\exp(\hat{\zeta}_g+\boldsymbol{m}_{\boldsymbol{c}}^{\boldsymbol{T}}\hat{\boldsymbol{\beta}}_g+\hat{\boldsymbol{w}}_{\boldsymbol{c}}^{\boldsymbol{T}}\hat{\boldsymbol{\alpha}}^*_g),\psi=\hat{\psi}^*_g,\pi^0=\hat{\pi}^0_{gc})\\ b_{gc} &= F(y_{gc}+1;\mu_{gc}=\exp(\hat{\zeta}_g+\boldsymbol{m}_{\boldsymbol{c}}^{\boldsymbol{T}}\hat{\boldsymbol{\beta}}_g+\hat{\boldsymbol{w}}_{\boldsymbol{c}}^{\boldsymbol{T}}\hat{\boldsymbol{\alpha}}^*_g),\psi=\hat{\psi}^*_g,\pi^0=\hat{\pi}^0_{gc})\end{aligned}$$

where $F(.)$ is the ZINB c.d.f, $F^{-1}(.)$ its inverse, $\boldsymbol{m}_c$ is the c$^{th}$ row of matrix $\boldsymbol{M}$, $\hat{\boldsymbol{w}}_{\boldsymbol{c}}$ the c$^{th}$ row of matrix $\hat{\boldsymbol{W}}$, $U(a,b)$ denoted a random variable uniformly distributed over the interval $(a,b)$, $\bar{\boldsymbol{w}}$ is the vector of the average level $N^{-1}\sum_{c=1}^N\hat{\boldsymbol{w}}_{\boldsymbol{c}}$ of unwanted variation and $\bar{\pi}^0_g=N^{-1}\sum_{c=1}^N\hat{\pi}^0_{gc}$ is the average zero-inflated parameter for gene $g$.

### Mean and variance of pseudo-cells

For the NB model the pseudo-cell count $z_{gj}$ comes from the pool-and-divide process with the following probability specification

$$z_{gj}\mid s_{gj}\;\sim\;\text{Binomial}(s_{gj},1/n_j).$$

Here $s_{gj}$ itself represents the sum of $n_j$ i.i.d negative binomial random variables. Further, let us assume that the cells that make up pool $j$ are sufficiently homogeneous that the expected value for the gene $g$ count is approximately $\mu_{gj}$ for all cells in the pool, and likewise its variance is approximately $\mu_{gj}+\psi_g\mu^2_{gj}$. It can then be shown that

$$\begin{aligned}E(s_{gj}) &\approx n_j\mu_{gj}\\ Var(s_{gj}) &\approx n_j\mu_{gj}+n_j\psi_g\mu^2_{gj}.\end{aligned}$$

Using rule of total expected value and total variance, we have

$$\begin{aligned}E(z_{gj}) &= E(E(z_{gj}\mid s_{gj}))\\ &= E\left(\frac{s_{gc}}{n_j}\right)\\ &= \mu_{gj}\end{aligned}$$

and

$$
\begin{aligned}
Var(z_{gj}) &= Var(E(z_{gj} \mid s_{gj})) + E(Var(z_{gj} \mid s_{gj})) \\
&= Var\left(\frac{s_{gc}}{n_j}\right) + E\left(\frac{s_{gj}(n_j - 1)}{n_j^2}\right) \\
&= \left(\frac{\mu_{gj} + \psi_g \mu_{gj}^2}{n_j}\right) + \left(\frac{\mu_{gj}(n_j - 1)}{n_j}\right) \\
&= \mu_{gc} + \frac{\psi_g}{n_j}\mu_{gj}^2
\end{aligned}
$$

Hence the pseudo-cell counts still have the quadratic mean-variance relationship of an NB random variable but with $\frac{\psi_g}{n_j}$ as dispersion parameter instead of $\psi_g$ .
